# Knockoff procedure improves causal gene identifications in conditional transcriptome-wide association studies

**DOI:** 10.1101/2025.02.05.636660

**Authors:** Xiangyu Zhang, Lijun Wang, Jia Zhao, Hongyu Zhao

## Abstract

Transcriptome-wide association studies (TWASs) have been developed to nominate candidate genes associated with complex traits by integrating genome-wide association studies (GWASs) with expression quantitative trait loci (eQTL) data. However, most existing TWAS methods evaluate the marginal association between a single gene and the trait of interest without accounting for other genes within the same genomic region or the same gene from different tissues. Additionally, false-positive gene-trait pairs can arise due to correlations with the direct effects of genetic variants. In this study, we introduce TWASKnockoff, a new knockoff-based framework for detecting causal gene-tissue pairs using GWAS summary statistics and eQTL data. Unlike marginal testing in traditional TWAS methods, TWASKnockoff examines the conditional independence for each gene-trait pair, considering both correlations in cis-predicted expression across genes and correlations between gene expression levels and genetic variants. TWASKnockoff estimates the theoretical correlation matrix for all genetic elements (cis-predicted expression across genes and genotypes for genetic variants) by averaging estimations from parametric bootstrap samples and then performs knockoff-based inference to detect causal gene-trait pairs while controlling the false discovery rate (FDR). Through empirical simulations and an application to type 2 diabetes (T2D) data, we demonstrate that TWASKnockoff achieves superior FDR control and improves the average power in detecting causal gene-trait pairs at a fixed FDR level.

## 1 Introduction

Genome-wide association studies (GWASs) have achieved remarkable success in detecting tens of thousands of genetic variants that are associated with complex traits [1]. However, the causal genes mediating the effects of genetic variants on the trait are rarely ascertainable by analyzing GWAS data alone [2]. This interpretation challenge has motivated the development of methods to prioritize causal genes at the GWAS loci.

Several methods have been proposed to leverage multi-omic data to nominate candidate genes from GWASs. One family of such methods is transcriptome-wide association studies (TWASs) [3, 4], which utilize expression reference panels (cohorts with gene expression and genotype data) to discover gene–trait associations from GWASs [5]. First, predictive models of expression variation are learned based on expression reference panels to predict gene expression for each individual in the GWAS cohort. However, most existing TWASs evaluate the marginal association between a single gene and the trait of interest without accounting for other genes within the same genomic region or the same gene from different tissues [5, 6]. The marginal testing approach does not account for the correlation among gene expressions within a genomic region, which can result in the identification of multiple gene associations in a localized area and an increased rate of false signals [7, 8]. Additionally, false-positive gene-tissue pairs can arise due to linkage disequilibrium (LD) between expression quantitative trait loci (eQTLs) and nearby genetic variants (e.g., single nucleotide polymorphisms, often called cis-SNPs), resulting in correlations between gene expressions and the direct effects of genetic variants [5, 9, 10].

To reduce false positives in marginal TWASs and prioritize the mapping of causal genes, several methods have been developed to jointly model a small number to a few dozen genes that reside in the same region [7]. Specifically, GIFT (Gene-based Integrative Fine-mapping through conditional TWAS) examines one genomic region at a time, jointly models the gene expression levels of all genes in the focal region, and performs TWAS conditional analysis in a maximum likelihood framework [11]. GIFT explicitly models the gene expression correlation and cis-SNP LD across different genes in the region and accounts for the uncertainty in the constructed gene expression in the GWAS cohort. However, GIFT assumes the absence of direct effect of genetic variants, which can lead to power loss and inflated false positives. Analysis of GWAS summary statistics for 42 traits and cis-eQTL summary statistics for 48 tissues from the Genotype-Tissue Expression (GTEx) suggested that averaging across traits, only 11±2% of heritability was mediated by assayed gene expression levels [9]. Consequently, GIFT’s assumption that the quantitative trait of interest is a linear combination of the expression levels of all genes in a local region may not be met in real-world data applications.

Model-X knockoffs is a broad and flexible variable selection framework in high-dimensional settings [12, 13], enabling practitioners to select variables that retain dependence with the response conditional on all other covariates while controlling the false discovery rate (FDR). Several knockoff-based methods have been developed for genetic applications, especially GWASs [14, 15]. Additionally, TWAS-GKF, a knockoff-based method, has been proposed to identify candidate trait-associated genes at the genome-wide level without accounting for non-mediated genetic variants [16]. However, the use of knockoff-based inference to jointly model candidate genes together with the direct effects of genetic variants within a local region remains unexplored.

We introduce TWASKnockoff, a new knockoff-based framework designed to identify causal genes for the trait of interest using GWAS summary statistics and eQTL data. TWASKnockoff extends the knockoff inference to identify disease-causing genes. Unlike the marginal testing performed in traditional TWAS methods, TWASKnockoff examines the conditional independence for each gene-trait pair, considering both correlations in cis-predicted expression across genes and correlations between gene expression levels and genotypes of genetic variants. TWASKnockoff begins by constructing gene expression prediction models using eQTL data, employing basic statistical methods such as elastic net and lasso regression. These models are used to predict summary statistics of gene expression in the GWAS cohort. Methodologically, we derive the correlation matrix for all genetic elements, including cis-predicted expression across genes and genotypes for genetic variants. TWASKnockoff approximates this theoretical correlation matrix by averaging estimations from parametric bootstrap samples. Finally, TWASKnockoff applies the GhostKnockoff procedure [17, 18] to select causal gene-trait pairs while maintaining FDR control.

We performed simulations and real data analyses for type 2 diabetes (T2D) to compare the performance of TWASKnockoff with GIFT in FDR control and power. TWASKnockoff outperformed GIFT by achieving superior FDR control and enhanced the average power for detecting genes causally associated with complex traits at a fixed FDR level. Besides, the improved estimation of the correlation matrix through bootstrap sampling further improved the performance of TWASKnockoff.

## 2 Methods

We consider a GWAS with *n*_1_ individuals and an eQTL study with *n*_2_ individuals. TWAS-Knockoff estimates the test statistics for genetic elements (including gene expression levels and non-mediated genetic variants) within a genomic region under a two-level model:

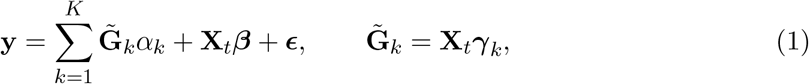

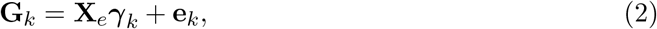

where **y** denotes the *n*_1_-vector of quantitive trait of interest, *K* denotes the total number of genes, **X**_*t*_ denotes the *n*_1_ × *p* matrix of genotypes for SNPs within this region in the GWAS cohort, **X**_*e*_ denotes the *n*_2_ × *p* matrix of genotypes for the same *p* cis-SNPs in the gene expression study, **G**_*k*_ is the *n*_2_-vector of the observed gene expression of gene *k*, 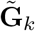 is the *n*_1_-vector of the predicted gene expression of gene *k, α*_*k*_ denotes the scalar effect of cis-gene expression in gene *k* on trait of interest, and ***γ***_*k*_ is the *p*-vector of causal cis-eQTL effect sizes of genetic variants on gene expression in gene *k*. Both ***ϵ*** and **e**_*k*_ are vectors of error terms following independent normal distributions. For each ***γ***_*k*_, we only consider the cis-SNPs for gene *k*, and set the effect sizes of other SNPs to zero. Without loss of generality, we assume **y, X**_*t*_, **X**_*e*_, and **G**_*k*_ have already been standardized.

Suppose **G** = [**G**_1_, …, **G**_*K*_] is the matrix of gene expression levels, ***α*** = [*α*_1_, …, *α*_*K*_]^*T*^ is the *K*-vector of the effects of gene expression levels on the trait of interest, ***γ*** = [***γ***_1_, …, ***γ***_*K*_] is the *p* × *K* matrix of cis-eQTL effect sizes of each variant on all gene expression levels. Thus we can rewrite the above model into:

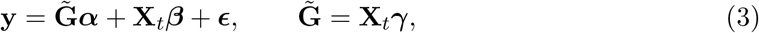

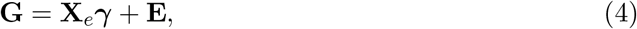

where **E** = [**e**_1_, …, **e**_*K*_] is the *n*_2_ × *K* matrix of random errors.

### 2.1 Effect size estimates for cis-eQTL

Based on **G** and **X**_*e*_, we can estimate the effect size ***γ***_*k*_ for each gene *k*, denoted as 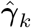. The gene expression level for gene *k* in the GWAS population can then be estimated as

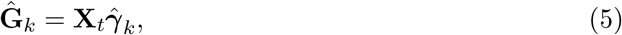

An accurate estimation 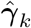 should produce 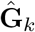 that closely approximates the true gene expression level for gene, i.e., 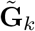. To estimate ***γ***_*k*_, we employed the elastic net regression method. However, in certain cases, the elastic net could not select any significant variables, resulting in noninformative predictions for 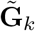. To address this limitation, we implement the following procedure to estimate ***γ***_*k*_ :

1. First, we fit the eQTL model (2) using the elastic net regression method. The penalty parameter is selected to minimize the cross-validation error.
2. If the elastic net model fails to identify any significant variables, we use ridge regression to fit the eQTL model, again selecting the penalty parameter to minimize the cross-validation error.

### 2.2 Improved correlation matrix estimate

Suppose 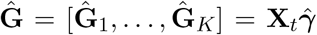 is the matrix of estimated gene expression levels in the GWAS cohort, where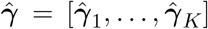. We want to solve the variable selection problem under model (1) using the knockoff framework, which requires an accurate estimation of the correlation structure for all genetic elements.

Let *G* = [*G*_1_, …, *G*_*K*_]^*T*^ represent the vector of imputed gene expression levels for *K* genes in GWAS, and *X*_*t*_ = [*X*_*t*,1_, …, *X*_*t,p*_]^*T*^ denote the random vector of standardized genotypes for *p* SNPs in GWAS. Therefore, we have 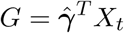. We define genetic elements *W* = [*G*^*T*^, *X*^*T*^]^*T*^, containing both gene expression levels and non-mediated genetic variants. To perform variable selection via knockoff, we need to model the covariance matrix of all the genetic elements:

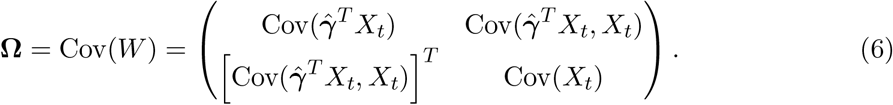

If the genotype matrix **X**_*t*_ is observable, we can directly estimate **Ω** by computing the empirical covariance matrix of 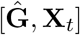. However, in most cases, instead of directly observing **X**_*t*_, we only have access to an estimate of the LD matrix, denoted as 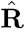, which is derived from an external reference panel. Here, **R** represents the true LD matrix. Under the scenario that 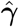 is fixed, **Ω** can be estimated as follows:

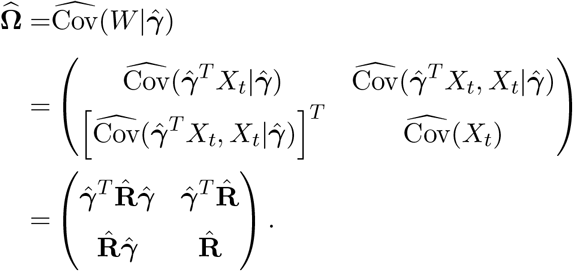

Although this empirical approach to calculating **Ω** is straightforward and intuitive, it overlooks the randomness introduced by the estimation of the size of the cis-eQTL effect,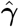. This simplification inevitably results in a biased estimation of both the covariance matrix among imputed gene expression levels and the covariance matrix between imputed gene expression levels and genotypes. To improve the accuracy of **Ω** estimation, it is important to recognize that 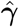 is a random matrix following a certain distribution. The covariance matrix of genetic elements can then be calculated by the law of total covariance:

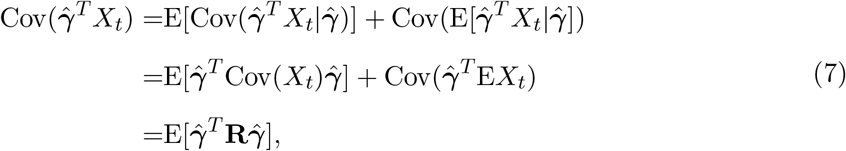

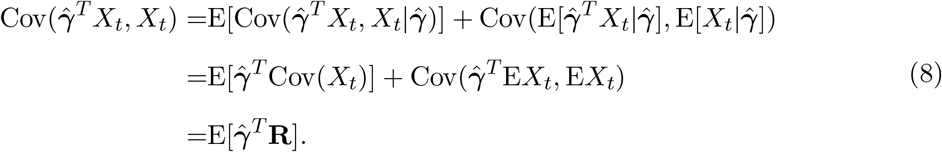

Assume that the coefficients estimated by the eQTL study are given by 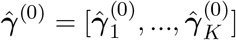. Using parametric bootstrap samples, we generate a total of *B* bootstrap estimates, denoted as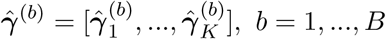, by resampling residuals:

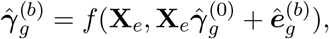

where 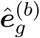 is randomly sampled from a normal distribution with mean zero and variance estimated by 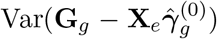. The function *f* denotes the method used to estimate the eQTL effect sizes based on observed data, such as elastic net or ridge regression.

Therefore, we can estimate (7) and (8) by:

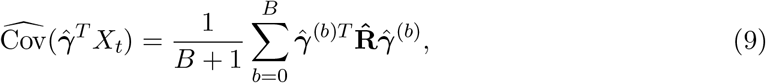

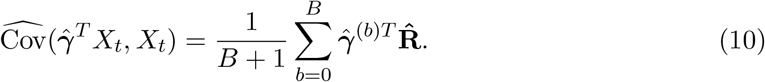

By substituting equations (9) and (10) into (6), we derive an improved estimation of **Ω**, referred to as the bootstrap estimation of the correlation matrix in this manuscript. We fixed the number of bootstrap samples *B* = 9 throughout this project. The effectiveness of the bootstrap estimation in increasing accuracy and stability for both gene-gene and gene-SNP correlations was demonstrated through simulations (see Supplementary materials for more details).

### 2.3 Variable selection via TWASKnockoff

We first give a brief review of model-X knockoffs and GhostKnockoff. Considering a very general conditional model where the response *y* can depend in an arbitrary fashion on the covariates *X*_1_, …, *X*_*p*_, and the observations (*X*_*i*1_, …, *X*_*ip*_, *y*_*i*_), *i* = 1, …, *n*, are independently and identically distributed. Model-X knockoffs aims to discover which variables are truly associated with the response while controlling the FDR.

To implement model-X knockoffs, the first step is to construct an *n* × *p* matrix 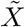 of knockoffs such that the following two conditions hold:

1. 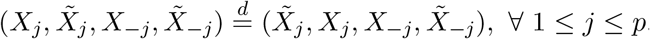.
2. 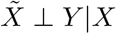.

In other words, for each feature *X*_*j*_, we construct a “knockoff” feature 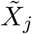 to imitate the correlation structure of the original features. After that, the feature statistic 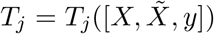 is computed for each pair of original and knockoff variables, and a large positive value of *T*_*j*_ provides evidence against the hypothesis that *X*_*j*_ is null.

To control the FDR below the desired level *q*, the knockoff filter selects the variables {*j* : *T*_*j*_ ≥ *T*}, where

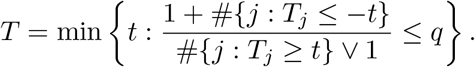

For each covariate *X*_*i*_ with feature statistic *T*_*i*_, the *q*-value is defined as

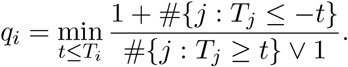

Selecting covariates at a target FDR level *q* is equivalent to selecting variants with *q*_*i*_ *< q*.

Model-X knockoffs requires individual-level data to perform variable selection. To address this limitation, GhostKnockoff modified the original knockoff framework to allow efficient knockoff-based inference using freely available GWAS summary statistics and LD information, which can directly generate the knockoff feature statistics per variant without the need to generate individual-level knockoffs.

The TWAS summary statistics z-scores for a specific gene *k* can be calculated using the GWAS summary statistic z-scores, the LD matrix, and the predicted effect sizes for cis-eQTLs:

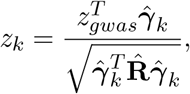

where *z*_*gwas*_ is the *p*-vector of GWAS summary statistic z-scores across variants in the risk region. By combining TWAS and GWAS summary statistics together with the correlation matrix of all genetic elements, we can perform variable selection under FDR control through the GhostKnockoff framework.

Under the model assumptions of TWASKnockoff, genetic variants can influence traits of interest through two distinct mechanisms. First, variants can act as cis-eQTLs for genes that are causal to the trait of interest. Second, variants can exert a direct effect on the trait without mediation through gene expression levels. To mitigate the power loss caused by conflating these two causal pathways, we exclude cis-eQTLs identified during gene expression mapping from the knockoff model of all genetic elements. Specifically, we remove significant genetic variants identified by the elastic net model in the gene expression prediction step. For genes where no significant variants are selected, we retain all cis-SNPs in the model.

## 3 Simulation

### 3.1 FDR control via TWASKnockoff

Effective control of the FDR often requires a sufficiently large number of discoveries. When the number of genes is adequate, variable selection and the calculation of *q*-values can be performed directly based on the feature statistics of the estimated gene expression levels, 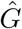. However, the number of candidate genes within a given risk region is often too small to enable direct FDR control. To address this limitation, we adopt an alternative approach that performs variable selection across all genetic elements, including both imputed gene expressions and cis-SNPs. This method controls the FDR of all genetic elements and then identifies the set of genes from the significant variants. The key idea is to leverage the larger set of genetic variants to facilitate effective FDR control for genes when the number of candidate genes is insufficient.

We conducted a simulation study to demonstrate that TWASKnockoff achieves effective FDR control by jointly modeling candidate genes and cis-SNPs within risk regions. We simulated both gene expression levels and quantitative phenotypes based on 1,000 and 20,000 unrelated European samples from the UK Biobank (UKBB), respectively. We randomly selected 100 risk regions on chromosome 1, each containing 100 adjacent and non-overlapping candidate genes. The proportion of causal genes contributing to the trait of interest was set at 20%. Each candidate gene contains 20 cis-SNPs from the common part of the UKBB and HapMap3 datasets. We assumed that there existed two eQTLs for each candidate gene, which explained 20% variation of the gene expression level. For simplicity, we excluded the direct effect of non-mediated SNPs. The total heritability explained by causal genes within the risk region was fixed at 0.1.

We compared the performance of two variable selection strategies in controlling the FDR within TWASKnockoff:

1. **Gene-based selection:** Identifying significant genes based solely on feature statistics of gene expression levels.
2. **Gene and SNP-based selection:** Identifying significant genes based on feature statistics of all genetic elements, including genes and cis-SNPs.

The performance of these strategies was assessed in terms of FDR control and power (Figure 1). We tested various feature statistics within TWASKnockoff, including lasso coefficients (`lasso`), lasso coefficients with approximated *λ* (`lasso.approx.lambda`), marginal empirical correlations (`marginal`), the element-wise square of z-scores (`squared.zscore`), and posterior inclusion probability produced by SuSiE [19] (`susie`). The correlation matrix of genetic elements was estimated using parametric bootstrap samples. Both strategies demonstrated effective FDR control relative to the *q*-value threshold. However, selecting significant genes solely based on feature statistics of estimated gene expression levels resulted in an FDR closely aligned with the *q*-value threshold. In contrast, the gene and SNP-based selection approach was more conservative, resulting in stricter FDR control. Additionally, feature statistics derived from lasso regression outperformed other methods, achieving a significantly lower FDR while maintaining comparable power.

**Figure 1:**
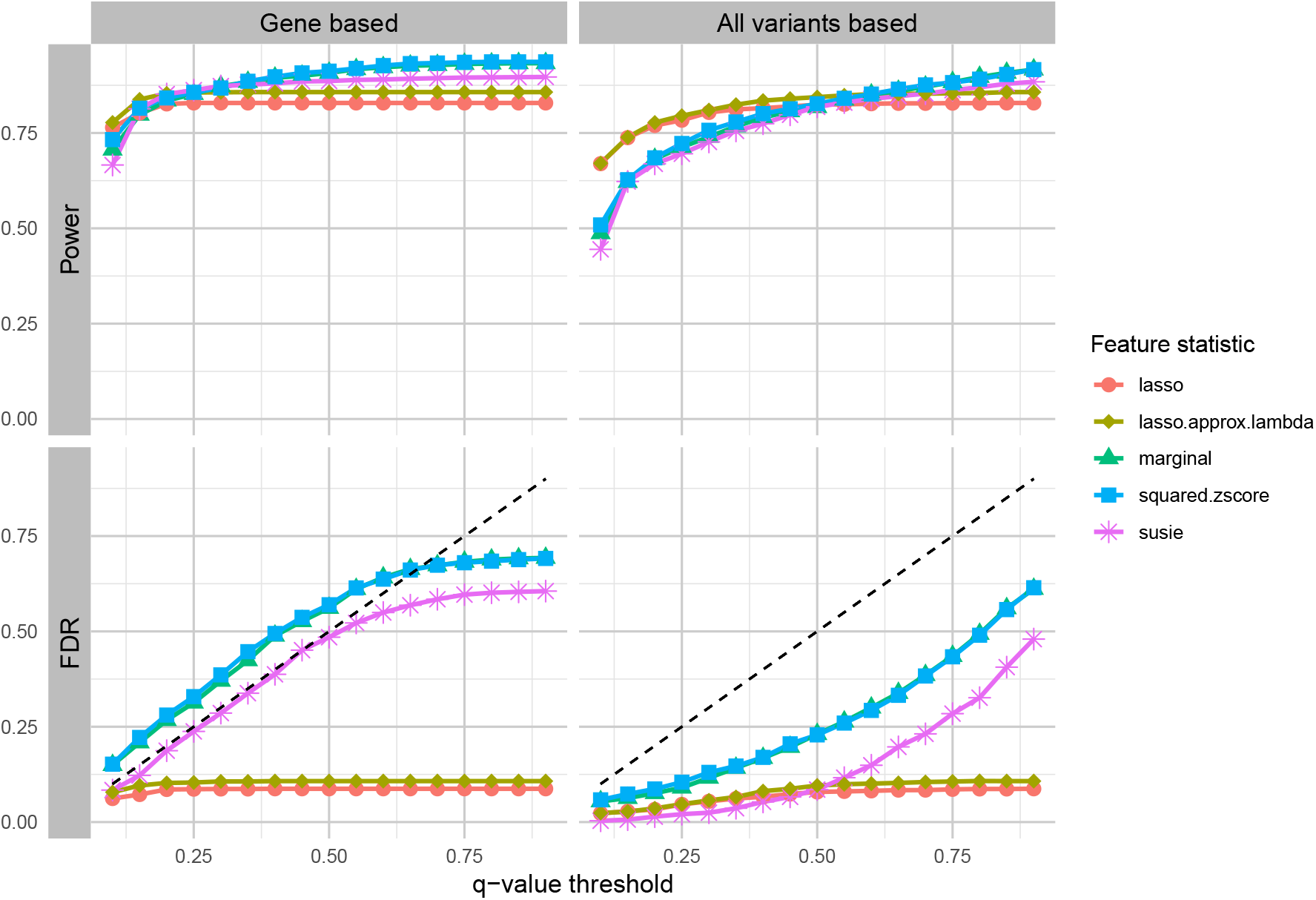
Comparison between two FDR control strategies. The FDR of genes was controlled within genes only (left) and all genetic elements (right) in TWASKnockoff, with the correlation matrix estimated with parametric bootstrap samples. In this simulation, all the heritability was explained by causal genes within the risk region, and we considered five feature statistics in the GhostKnockoff framework.

The same simulation was also performed using the empirical estimation of the correlation matrix (Figure S**??**). The results further validated that variable selection based on all genetic elements effectively controlled the FDR. The degree of conservativeness in FDR control was largely influenced by the comparability between the proportion of causal elements within the gene set and those within the SNP set (see Supplementary Materials for additional details).

### 3.2 Comparison of TWASKnockoff with GIFT

We conducted extensive simulations to evaluate the performance of TWASKnockoff and compare it with GIFT. These simulations used imputed genotype data from the UKBB and involved simulated gene expression levels and quantitative phenotypes based on 1,000 and 50,000 unrelated participants of white British ancestry, respectively. We randomly selected 100 risk regions on chromosome 1, each containing *K* adjacent but non-overlapping candidate genes. Each candidate gene contained 100 HapMap3 cis-SNPs, of which two were selected as eQTLs, explaining 20% of the variation in gene expression level. The causal status of each candidate gene was simulated using a binomial distribution *B*(*K, P*_*e*_), where the success probability *P*_*e*_ was fixed at 0.2. To ensure that each risk region included at least one causal gene, if the binomial distribution produced no causal genes, the first gene was set to be causal. Gene expression levels and phenotypes were simulated following the two-level model described below:

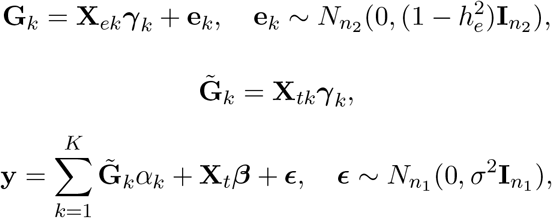

where **y** is the quantitative trait of interest, **X**_*t*_ is an *n*_1_ × *p* matrix of standardized genotypes for the *p* SNPs from the risk region in the GWAS, **X**_*ek*_ is an *n*_2_ × *p*_*k*_ matrix of standardized genotypes for the same *p*_*k*_ cis-SNPs of the *k*-th candidate gene in the eQTL study, and **X**_*tk*_ is an *n*_1_ × *p*_*k*_ matrix of standardized genotypes for the same *p*_*k*_ cis-SNPs of the *k*-th candidate gene in the GWAS cohort. We denote *h*_*e*_ as the variation explained by cis-eQTL for gene expression levels, **G**_*k*_ is the gene expression level for the *k*-th candidate gene in the eQTL study, and 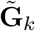 is the gene expression level for the *k*-th candidate gene in the GWAS cohort. We denote ***γ***_*k*_ as the eQTL effect size, which has non-zero elements only at *n*_*e*_ eQTLs. For each eQTL, we sampled the effect size from a normal distribution 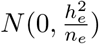.We assumed that the probability for any SNP within the given risk region to have a non-mediated effect on the trait of interest is *P*_*t*_. The total number of non-mediated SNPs, *n*_*nmv*_, was randomly sampled from a Poisson distribution *Pois*(*p*_0_*P*_*t*_), where *p*_0_ is the total number of genetic variants that are not eQTLs. For each non-mediated SNP *j*, the effect size *β*_*j*_ was sampled from a normal distribution 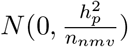. For all other SNPs, the non-mediated effect size was set to zero.

Additionally, we assigned *α*_*k*_ to be a fixed constant *c* if the *k*-th gene was causal and *α*_*k*_ = 0 otherwise.

To evaluate the performance of TWASKnockoff under different parameter settings, we considered two key parameters. The first parameter is the total proportion of phenotypic variance explained by genetic elements in the candidate region, denoted as *h*^2^. The second parameter is the proportion of heritability attributable to gene expression, denoted as *r*_*e*_. Assuming the total number of causal genes is *K*_1_, we solved the following equations to determine the values of 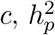, and *σ*^2^ in order to control *h*^2^ and *r*_*e*_ at the desired levels:

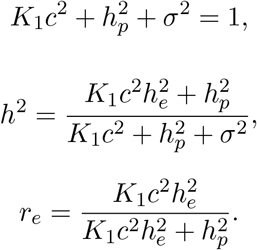

Solving these equations, we obtain:

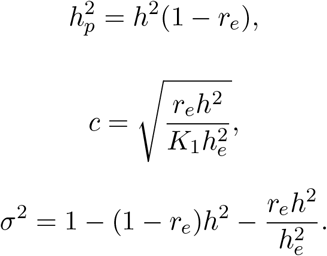

We performed simulations for each combination of *h*^2^ ∈ {1%, 2%, 4%} and *r*_*e*_ ∈ {20%, 50%, 100%}. The default settings for other parameters were as follows: *n*_1_ = 50, 000, *n*_2_ = 1, 000, *K* = 10, 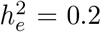, *n*_*e*_ = 2, *P*_*t*_ = 0.01, *P*_*e*_ = 0.2, *p*_*k*_ = 100 for *k* = 1, …, *K*. Notably, when *r*_*e*_ = 100%, there were no non-mediated effects, aligning with the model assumptions of GIFT.

#### 3.2.1 Simulation with the in-sample LD matrix

We first assessed the performance of TWASKnockoff and GIFT when the in-sample LD matrix was available. For TWASKnockoff, we utilized summary statistics and the in-sample LD matrix, considering two representative feature statistics: `lasso` and `squared.zscore`. To ensure a fair comparison, we used individual-level data as input for GIFT.

The performance of TWASKnockoff and GIFT was evaluated in terms of statistical power and FDR control when genetic elements in the region explained 1% of phenotypic variance (Figure 2). When *r*_*e*_ = 1, meaning all heritability was explained through gene expression levels, the model assumptions of GIFT were satisfied. Under this scenario, GIFT effectively controlled the FDR at the desired level and achieved high power. In contrast, TWASKnockoff exhibited a more conservative behavior, attributable to its inclusion of non-mediated genetic variants for FDR control.

**Figure 2:**
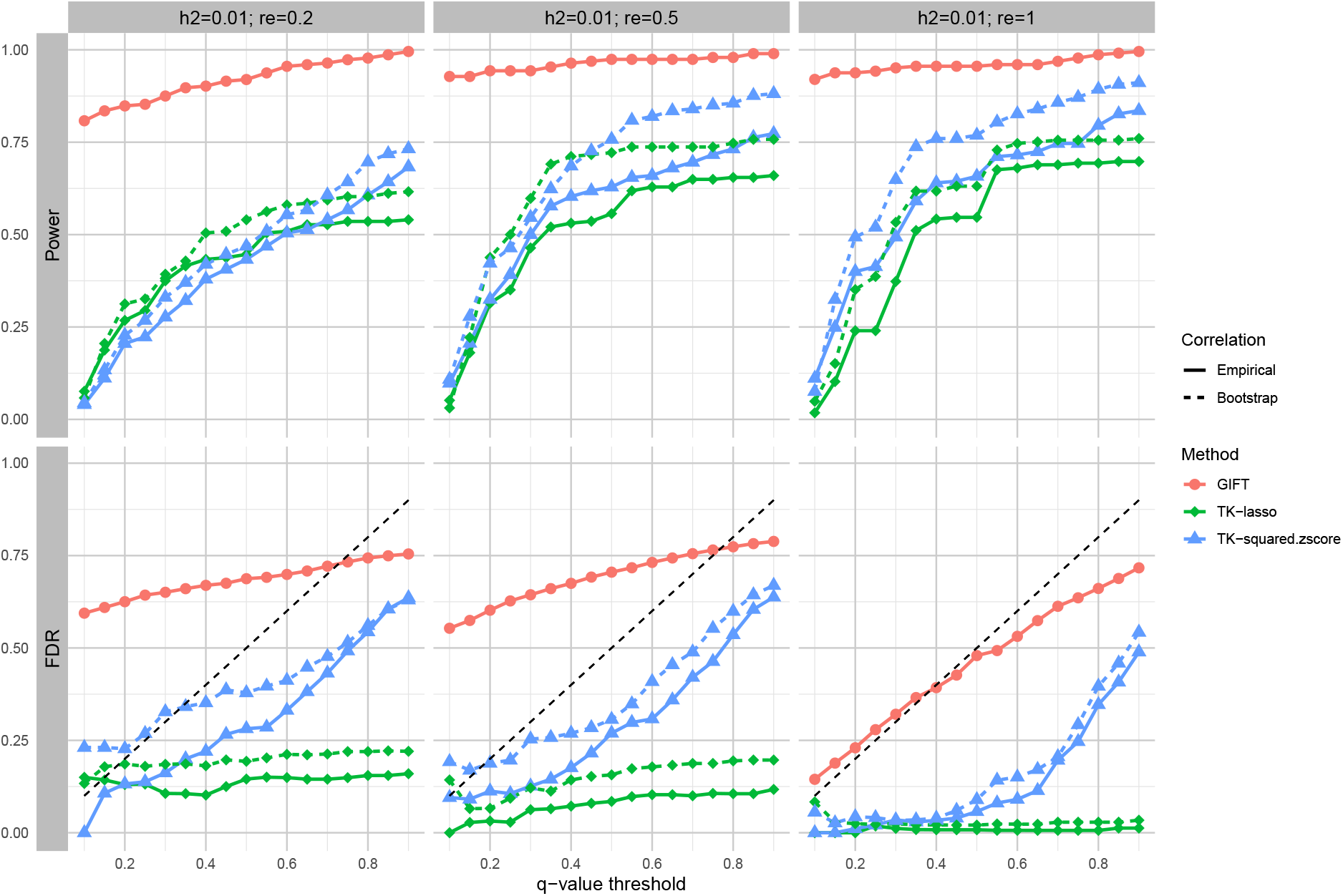
Comparison of TWASKnockoff and GIFT in simulated data with the in-sample LD matrix when *h*^2^ = 0.01. We assess the performance of GIFT and TWAS-Knockoff with two feature statistics: lasso coefficients (TK-lasso) and squared z-scores (TK-squared.zscore). For each feature statistic, we consider two estimation methods (empirical estimation and bootstrap samples) of the correlation matrix of genetic elements. The *q*-value threshold is selected from 0.1 to 0.9, with the black dashed line indicating the theoretical *q*-value level. For GIFT, we applied the Benjamini-Hochberg (BH) correction to perform FDR control.

Although GIFT performed well under the extreme condition where all heritability was mediated through gene expression (*r*_*e*_ = 1), it is important to recognize that, in most realistic scenarios, a substantial portion of heritability is not mediated by gene expression. When non-mediated effects of genetic variants were present (*r*_*e*_ = 20% or *r*_*e*_ = 50%), GIFT failed to control false discoveries, exhibiting an inflated FDR of up to 60% even under the strictest *q*-value thresholds. These uncontrolled false discoveries naturally led to a higher power, which can be explained as follows: GIFT’s inability to account for direct genetic effects resulted in false discoveries when non-mediated causal SNPs and the eQTLs of non-causal genes were located in the same LD block. Additionally, non-mediated causal SNPs within the cis-region of causal genes amplified the signal, increasing the likelihood of detection. In contrast, TWAS-Knockoff successfully controlled the FDR below the threshold by accounting for both the mediated effects of gene expression and the direct effects of non-mediated genetic variants. For both feature statistics (`lasso` and `squared.zscore`), the empirically estimated correlation matrix produced more conservative results, with lower FDR and reduced power. Notably, for TWASKnockoff using lasso, as the *q*-value threshold increased, both power and FDR stabilized rather than continuing to grow. This phenomenon arises from lasso’s sparse variable selection property: for non-significant variables, the knockoff method with lasso tends to assign feature statistics of zero. Consequently, these non-significant variables are excluded from consideration regardless of the *q*-value threshold applied. Similar patterns were observed across various simulation settings with different total heritabilities (*h*^2^), as shown in the Supplementary Figures.

To further compare the performance of TWASKnockoff and GIFT in identifying causal genes, we varied the threshold of test statistics for different methods (feature statistics for TWASKnockoff and *p*-values for GIFT) and evaluated the statistical power while controlling the FDR at a fixed level (Figure 3 A) when 50% of the heritability was explained by gene expression. TWASKnockoff consistently achieved a higher statistical power compared to GIFT. Moreover, estimating the correlation matrix via bootstrap sampling further enhanced the power of TWASKnockoff compared to using the empirical correlation matrix.

**Figure 3:**
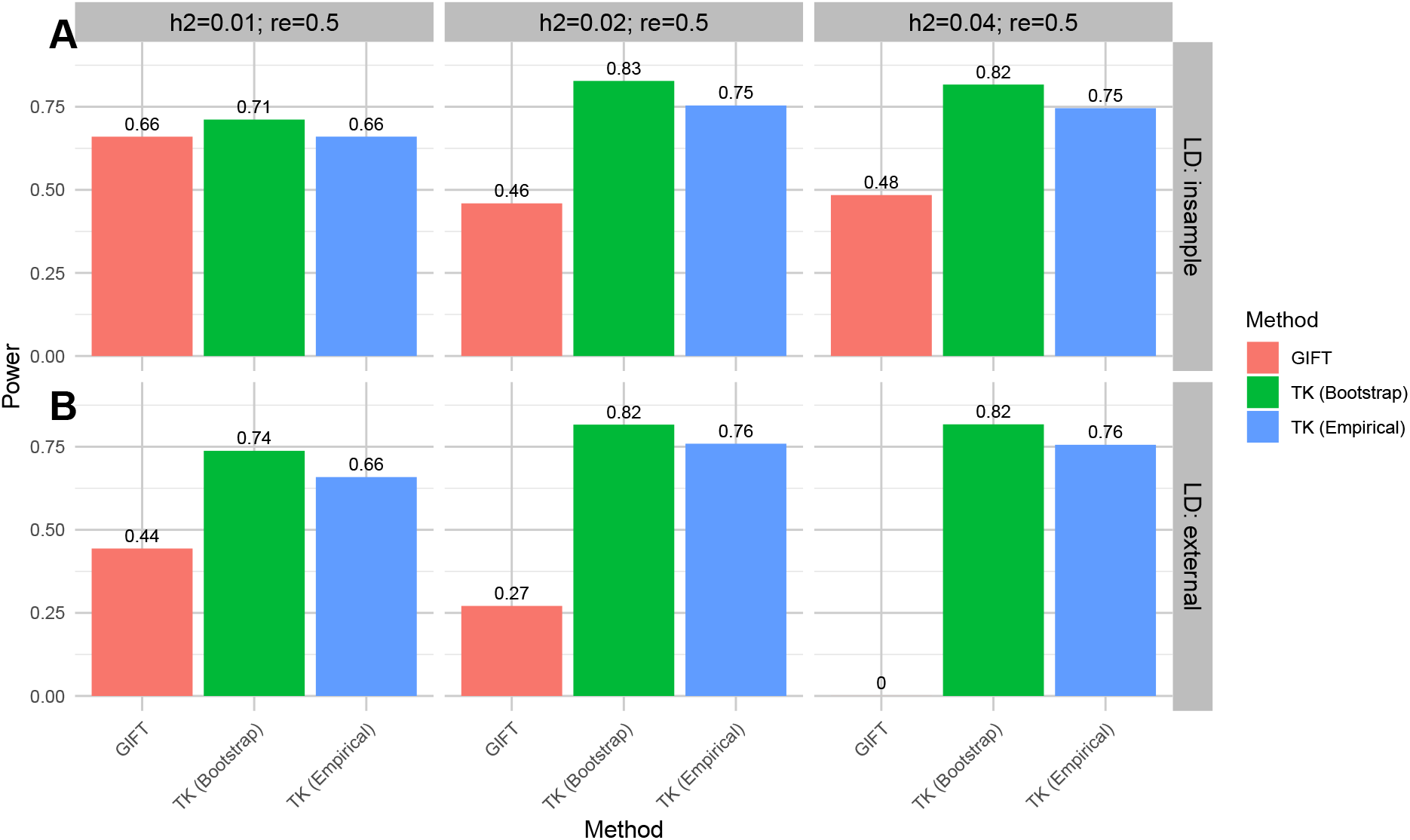
Comparison of TWASKnockoff and GIFT in power at fixed FDR = 0.2 with the in-sample LD matrix (A) and the external reference panel (B). We assess the performance of GIFT and TWASKnockoff with the feature statistic squared z-score (TK). For TWASKnockoff, we consider two estimation methods (empirical estimation and bootstrap samples) of the correlation matrix of genetic elements. We vary the total heritability (*h*^2^), with the proportion of heritability explained by gene expression (*r*_*e*_) fixed at 0.5.

#### 3.2.2 Simulation with the 1KG dataset as reference panel

To enhance the realism of the simulation settings, we also evaluated the performance of TWASKnockoff and GIFT using an external reference panel from the 1000 Genomes (1KG) Phase 3 dataset [20] (details provided in the Supplementary Materials). For GIFT, we utilized the summary statistics-based version (GIFT-ss), which takes summary statistics and LD matrices as input. Simulations were conducted to compare TWASKnockoff with GIFT in terms of statistical power and FDR control when the LD structure was inaccurate (Figure 4). When the reference panel provided an inaccurate LD structure, GIFT failed to maintain FDR control even when the simulation setting satisfied the model assumption of GIFT (*r*_*e*_ = 1). In contrast, TWASKnockoff demonstrated superior FDR control under these conditions. Notably, TWASKnockoff based on squared z-scores exhibited greater stability with respect to inaccuracies in the external reference panel compared to lasso-based feature statistics. We further extended these simulations across a range of settings with varying total heritabilities (*h*^2^) (see Supplementary Figures). As total heritability increased, both GIFT and lasso-based TWASKnockoff exhibited higher FDR along with increased power.

**Figure 4:**
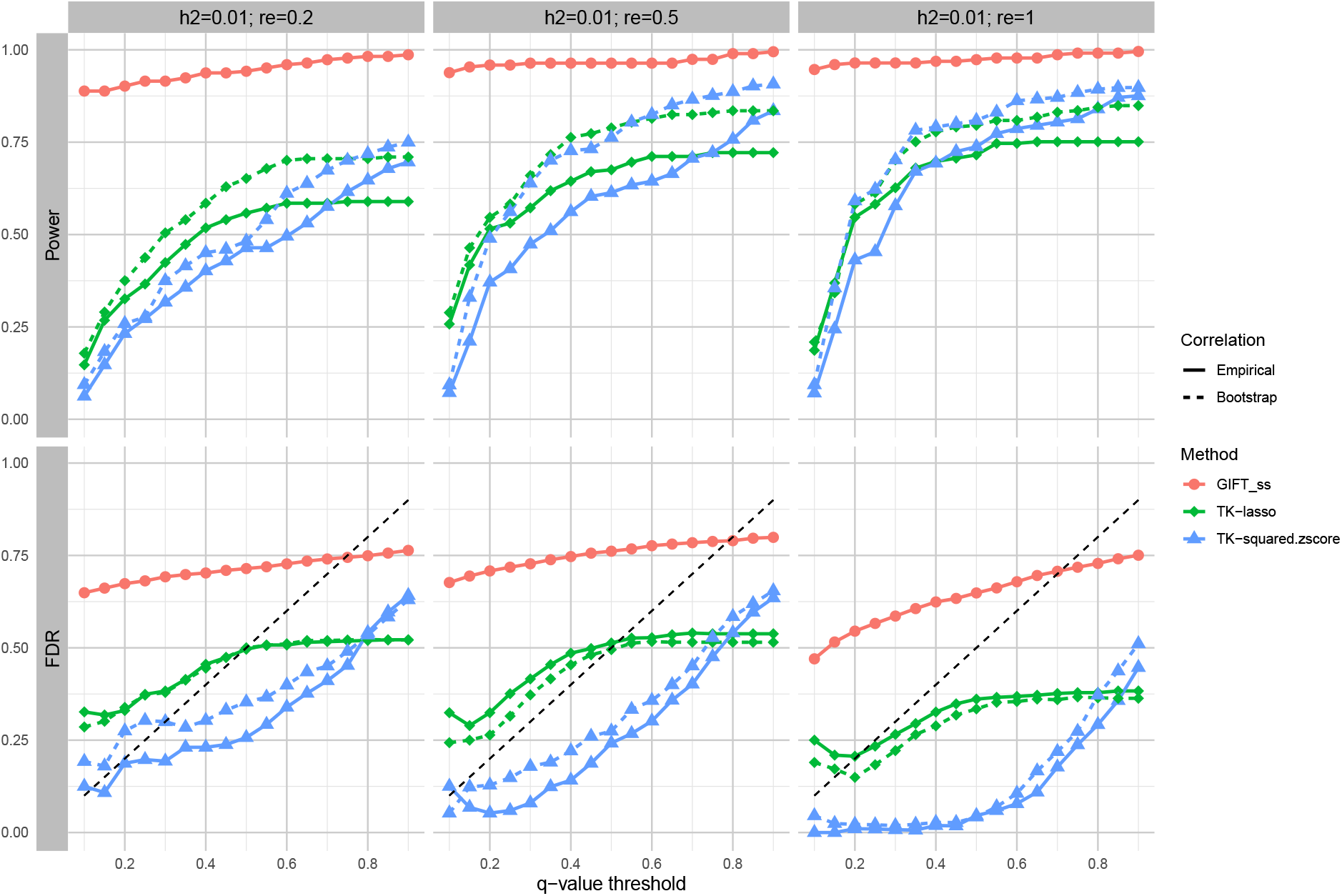
Comparison of TWASKnockoff and GIFT in simulated data with the external reference panel when *h*^2^ = 0.01. We assess the performance of GIFT and TWASKnockoff with two feature statistics: lasso coefficients (TK-lasso) and squared z-scores (TK-squared.zscore). For each feature statistic, we consider two estimation methods (empirical estimation and bootstrap samples) of the correlation matrix of genetic elements. The *q*-value threshold is selected from 0.1 to 0.9, with the black dashed line indicating the theoretical *q*-value level. For GIFT, we applied the Benjamini-Hochberg (BH) correction to perform FDR control.

We also evaluated the power of TWASKnockoff, with squared z-scores as the feature statistics, in comparison to GIFT at a fixed FDR of 0.2 (Figure 3 B). GIFT exhibited a substantial reduction in power when the in-sample LD matrix was replaced by an external reference panel. Furthermore, GIFT’s power dropped to zero in scenarios with a high total heritability, indicating its inability to control the FDR below 0.2, irrespective of the chosen *q*-value threshold.

In contrast, TWASKnockoff demonstrated remarkable stability when using the external reference panel. This robustness can be attributed to TWASKnockoff’s use of a regularized LD matrix for both in-sample LD and external reference panels, ensuring that the correlation matrix for all genetic elements remained positive-definite. Specifically, TWASKnockoff applied the following transformation to 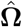:

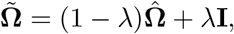

where *λ* was set to 0.1. While this transformation could slightly reduce the accuracy of the correlation structure—particularly for in-sample LD cases where the LD information is highly accurate, replacing 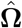 with 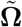 when applying to TWASKnockoff ensured the positive-definite property of correlation matrix and improved the stability of the algorithm.

Additionally, while TWASKnockoff accounted for non-mediated genetic variants alongside the candidate genes, it still demonstrated superior computational efficiency compared to GIFT. GIFT employs a parameter expansion version of the expectation-maximization (PX-EM) algorithm to maximize the joint likelihood, which is computationally intensive. As shown in Figure **??**, TWASKnockoff required less computational time than GIFT, even when the number of iterations for GIFT was set to 100, as recommended by GIFT v2.0. If the iteration parameter is increased to 1,000, as suggested by GIFT v1.0, the computational time for GIFT increases substantially (Figure **??**). Although the bootstrap-based estimation of the correlation matrix increased TWASKnockoff’s computational time, it remained faster than GIFT, regardless of whether GIFT utilized individual-level data or summary statistics. Besides, this bootstrap-based estimation step can be parallelized to further speed up the algorithm, which is our future work.

## 4 Real data applications

We conducted a conditional TWAS analysis on type 2 diabetes (T2D) by integrating gene expression datasets from ten T2D-relevant tissues from the Genotype-Tissue Expression (GTEx) Project v8 [21] with GWAS EUR ancestry meta-analysis summary statistics [22] (see Supplementary materials for details). The GTEx dataset provided tissue-specific gene expression levels and whole-genome sequencing data. The reference panel consisted of 503 unrelated European samples from the 1KG dataset. Based on the T2D summary statistics, we defined 484 candidate regions. These candidate regions were constructed around significant genetic association signals (*p*-value *<* 5 × 10^−10^), with the property that all SNPs within 250 Kb upstream and downstream of each signal belonged to the same candidate region. Overlapping regions were merged until no further overlap remained. For interpretability, we restricted our analysis to protein-coding genes with available gene expression data in the GTEx dataset.

We compared the performance of TWASKnockoff with GIFT in identifying causal genes for T2D while controlling the FDR using gene expression data from the ten T2D-related tissues. For TWASKnockoff, we selected significant genes with *q*-values below 0.1. For GIFT, *p*-values were adjusted using the Benjamini-Hochberg (BH) procedure, and genes with adjusted *p*-values less than 0.1 were identified as significant. Among 48,989 candidate gene-tissue pairs, TWASKnockoff identified 359 pairs (corresponding to 240 unique genes), while GIFT identified 11,879 gene-tissue pairs (corresponding to 2,135 unique genes) (Table 1).

**Table 1:**
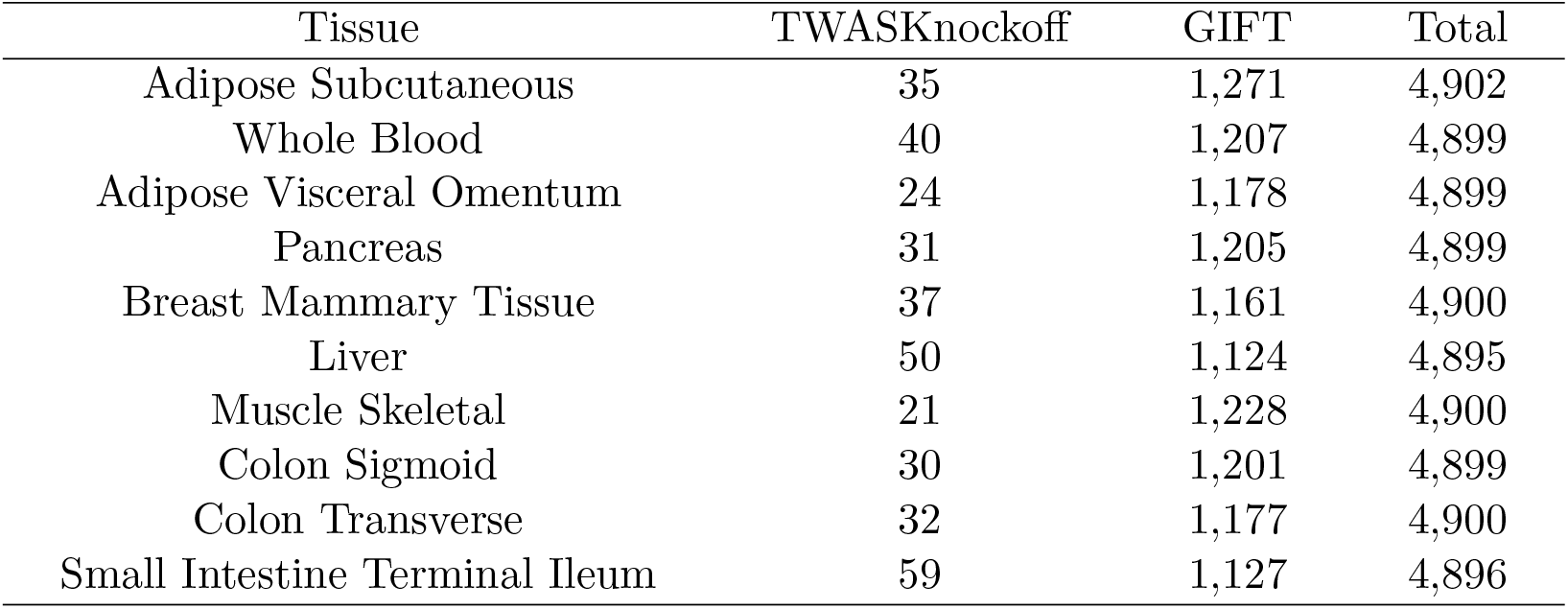
Summary of significant genes identified by TWASKnockoff and GIFT, and the total number of candidate genes for each GTEx tissue.

To validate the identified genes, we compared our results with T2D-related genes listed in GeneCards [23], a searchable, integrative database that provides comprehensive information on all human genes. GeneCards assigns a relevance score to each risk gene, indicating the strength of its association with the trait of interest. This score is calculated using Elastic-search (https://www.elastic.co/elasticsearch), which employs the Boolean model for document matching and the practical scoring function for relevance computation. For each GTEx tissue analyzed, we calculated the baseline relevance score for all candidate genes and the averaged relevance score of significant genes identified by TWASKnockoff and GIFT with T2D (Figure 5). Across all T2D-related tissues, genes identified by both TWASKnockoff and GIFT had higher average relevance scores compared to the baseline. However, the genes identified by TWASKnockoff consistently exhibited significantly higher relevance scores than those identified by GIFT, suggesting that TWASKnockoff identified genes that were more strongly associated with T2D. Notably, TWASKnockoff showed the most significant enrichment of relevance scores in tissues closely related to T2D, such as subcutaneous adipose (where the highest improvement was observed), whole blood, skeletal muscle, and pancreas.

**Figure 5:**
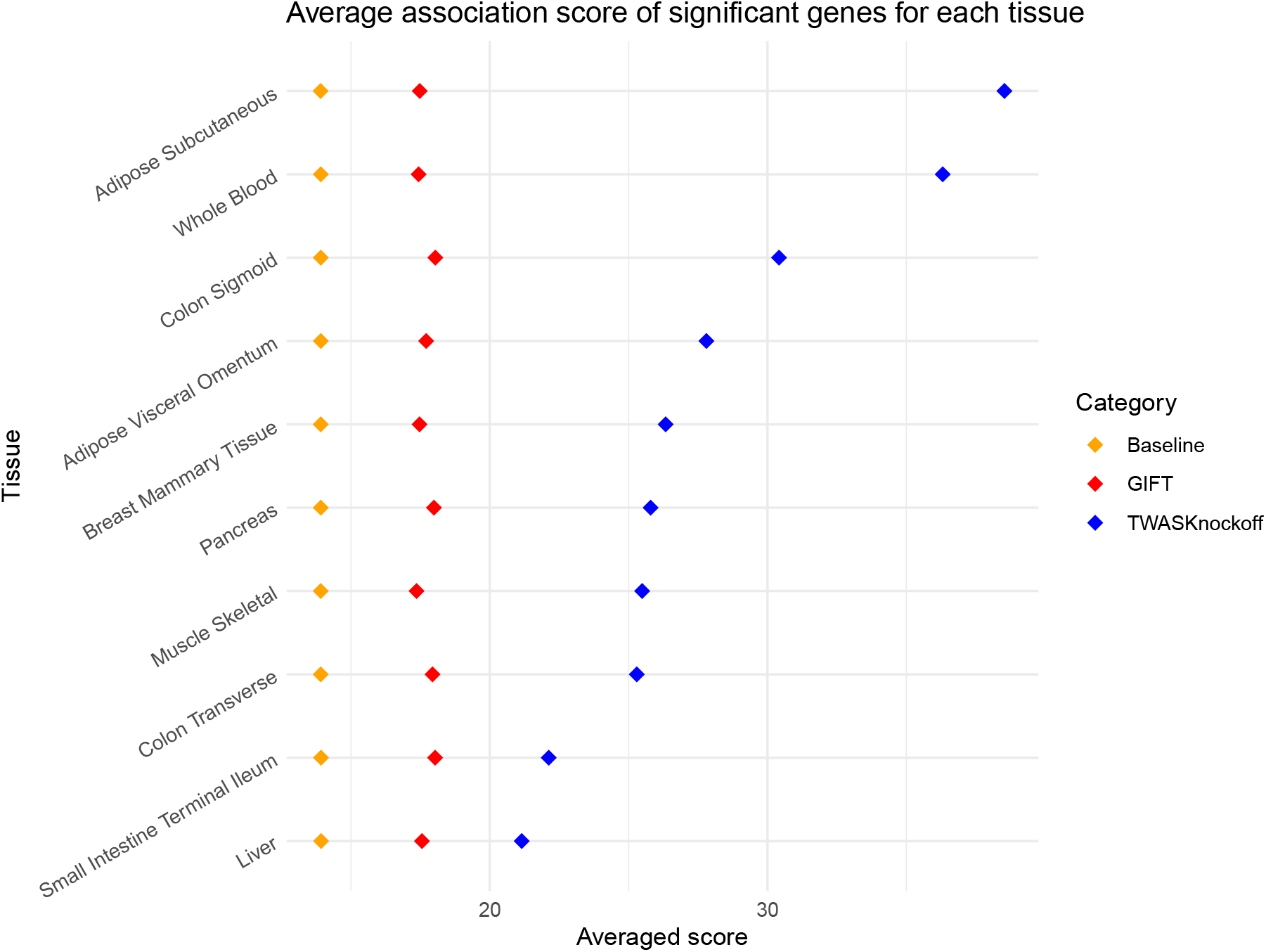
Tissue-specific averaged relevance scores for all candidate genes (baseline) and significant genes identified by TWASKnockoff and GIFT.

We highlight the pancreas (362 GTEx samples) as an example tissue closely linked to T2D. TWASKnockoff identified 31 significant genes with *q*-values less than 0.1 in the pancreas tissue (Table 2), 28 of which have been associated with T2D according to GeneCards. Several significant genes were strongly implicated in T2D susceptibility across multiple studies. For instance, TWASKnockoff identified TCF7L2, the most potent locus for T2D risk. The association of TCF7L2 with T2D has been consistently replicated in multiple populations with diverse genetic origins [24, 25]. Risk alleles of TCF7L2 are associated with reduced insulin secretion due to impaired beta cell function [26]. TWASKnockoff also identified Tumor Necrosis Factor-alpha (TNF-*α*), which is a pro-inflammatory cytokine that plays a critical role in the development of insulin resistance and the pathogenesis of T2D [27, 28]. In addition, several significant genes identified by TWASKnockoff have been recognized as key regulators for pancreatic development and, consequently, the pathogenesis of T2D. For example, Cyclin D2 (CCND2) plays a key role in regulating the transition of *β* cells from quiescence to replication and is essential for postnatal pancreatic *β* cell growth [29, 30, 31]. Moreover, overexpression of IGF2BP2 was found to disrupt pancreatic development, impair *β*-cell repair, and promote the translation of insulin-like growth factor 2 (IGF2) [32, 33, 34]. IGF2 itself serves as a key paracrine regulator of pancreatic growth and function, with its deletion leading to acinar and *β*-cell hypoplasia, postnatal growth restriction, and maternal glucose intolerance during pregnancy [35]. These findings demonstrate TWASKnockoff’s capability to identify disease-causing genes that are functionally relevant to the target tissue.

**Table 2:**
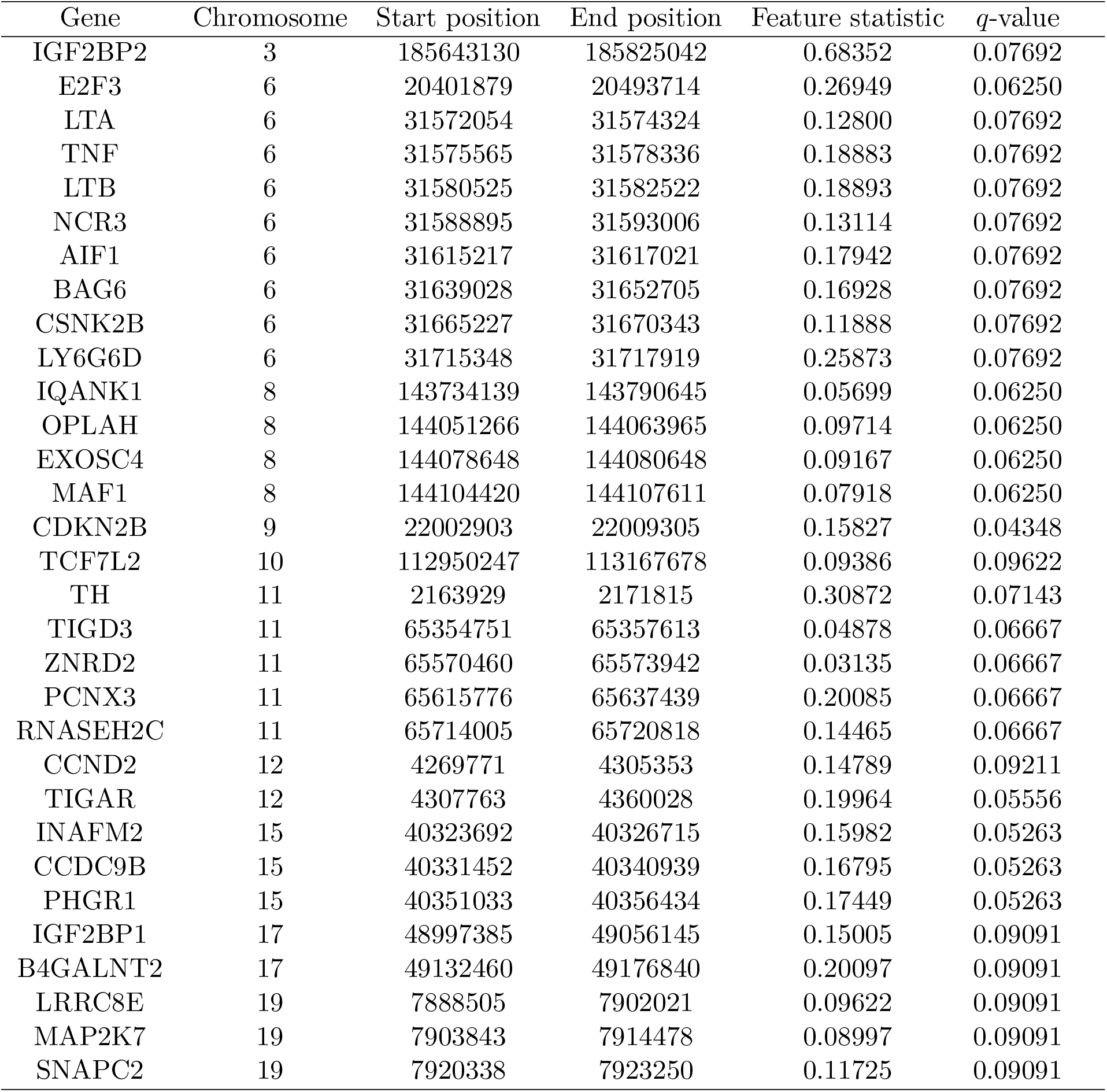
Summary of significant genes identified by TWASKnockoff with GTEx pancreas data in the T2D study.

| Gene | Chromosome | Start position | End position | Feature statistic | $q$-value |
| --- | --- | --- | --- | --- | --- |
| IGF2BP2 | 3 | 185643130 | 185825042 | 0.68352 | 0.07692 |
| E2F3 | 6 | 20401879 | 20493714 | 0.26949 | 0.06250 |
| LTA | 6 | 31572054 | 31574324 | 0.12800 | 0.07692 |
| TNF | 6 | 31575565 | 31578336 | 0.18883 | 0.07692 |
| LTB | 6 | 31580525 | 31582522 | 0.18893 | 0.07692 |
| NCR3 | 6 | 31588895 | 31593006 | 0.13114 | 0.07692 |
| AIF1 | 6 | 31615217 | 31617021 | 0.17942 | 0.07692 |
| BAG6 | 6 | 31639028 | 31652705 | 0.16928 | 0.07692 |
| CSNK2B | 6 | 31665227 | 31670343 | 0.11888 | 0.07692 |
| LY6G6D | 6 | 31715348 | 31717919 | 0.25873 | 0.07692 |
| IQANK1 | 8 | 143734139 | 143790645 | 0.05699 | 0.06250 |
| OPLAH | 8 | 144051266 | 144063965 | 0.09714 | 0.06250 |
| EXOSC4 | 8 | 144078648 | 144080648 | 0.09167 | 0.06250 |
| MAF1 | 8 | 144104420 | 144107611 | 0.07918 | 0.06250 |
| CDKN2B | 9 | 22002903 | 22009305 | 0.15827 | 0.04348 |
| TCF7L2 | 10 | 112950247 | 113167678 | 0.09386 | 0.09622 |
| TH | 11 | 2163929 | 2171815 | 0.30872 | 0.07143 |
| TIGD3 | 11 | 65354751 | 65357613 | 0.04878 | 0.06667 |
| ZNRD2 | 11 | 65570460 | 65573942 | 0.03135 | 0.06667 |
| PCNX3 | 11 | 65615776 | 65637439 | 0.20085 | 0.06667 |
| RNASEH2C | 11 | 65714005 | 65720818 | 0.14465 | 0.06667 |
| CCND2 | 12 | 4269771 | 4305353 | 0.14789 | 0.09211 |
| TIGAR | 12 | 4307763 | 4360028 | 0.19964 | 0.05556 |
| INAFM2 | 15 | 40323692 | 40326715 | 0.15982 | 0.05263 |
| CCDC9B | 15 | 40331452 | 40340939 | 0.16795 | 0.05263 |
| PHGR1 | 15 | 40351033 | 40356434 | 0.17449 | 0.05263 |
| IGF2BP1 | 17 | 48997385 | 49056145 | 0.15005 | 0.09091 |
| B4GALNT2 | 17 | 49132460 | 49176840 | 0.20097 | 0.09091 |
| LRRC8E | 19 | 7888505 | 7902021 | 0.09622 | 0.09091 |
| MAP2K7 | 19 | 7903843 | 7914478 | 0.08997 | 0.09091 |
| SNAPC2 | 19 | 7920338 | 7923250 | 0.11725 | 0.09091 |

To further illustrate the ability of TWASKnockoff to effectively control the FDR, we compared its performance with GIFT in the risk region surrounding TCF7L2, the most potent locus for T2D risk (Figure 6). This risk region includes six candidate genes with moderate gene-gene correlations. Table 3 summarizes the feature statistics and *q*-values estimated by TWASKnockoff for each candidate gene. Similarly, for GIFT, we report the causal effect estimates, *p*-values, and *q*-values adjusted by the BH procedure. Using a *q*-value threshold of 0.1, TWASKnockoff identified only TCF7L2 as a significant gene in this region. In contrast, GIFT yielded *q*-values below 0.1 for all candidate genes in the region. While other T2D-related genes may exist in this region, the very small *q*-values reported by GIFT indicate the limitation of this *p*-value-based method to control false discoveries effectively. These results demonstrate TWASKnockoff’s robustness in maintaining FDR control even in high-risk genomic regions.

**Table 3:**
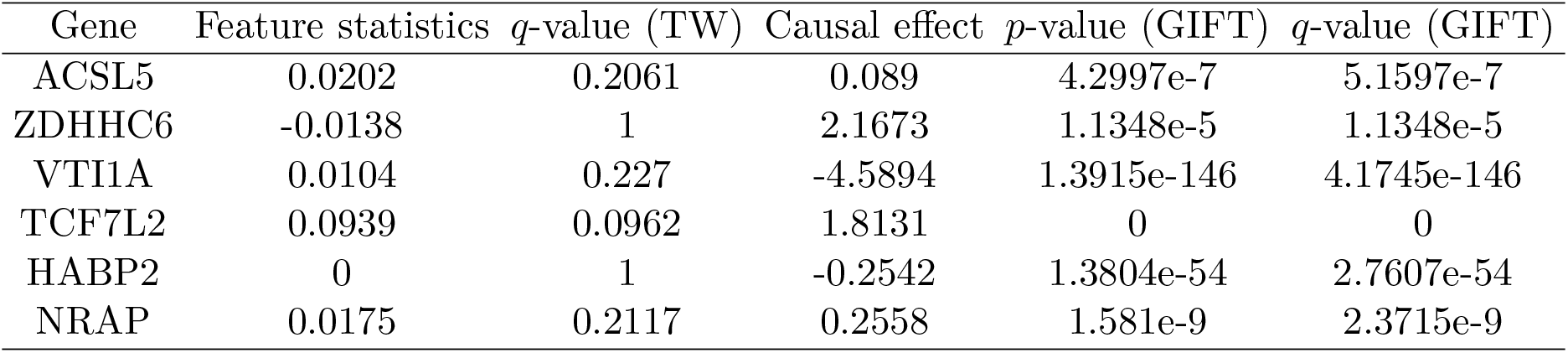
Comparison of TWASKnockoff and GIFT for candidate genes in the example risk region around TCF7L2.

**Figure 6:**
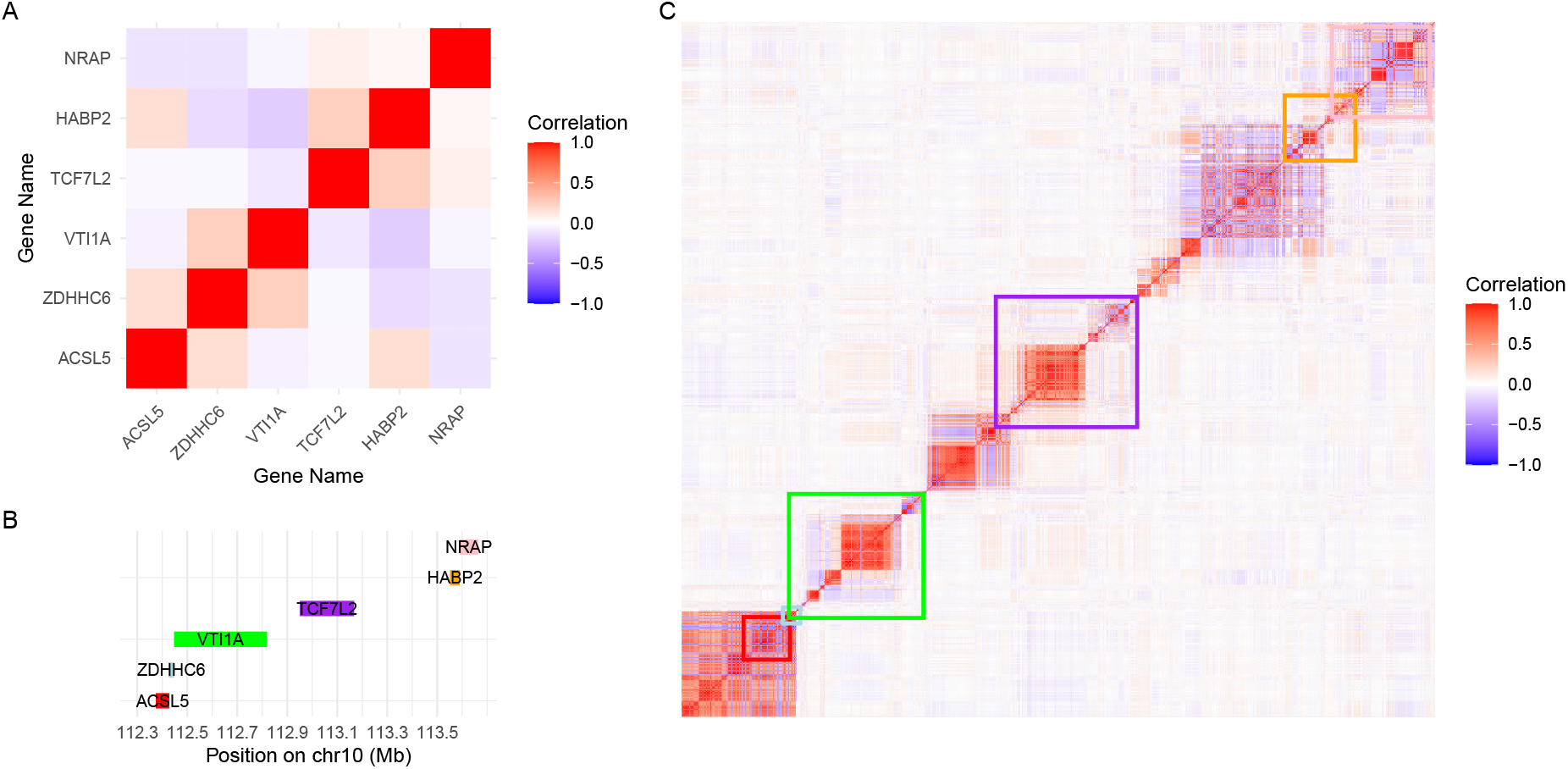
Information of the risk region around TCF7L2. We summarize the basic information for the example region including 6 candidate genes on Chromosome 10, including the correlation matrix of the gene expression level corresponding to each candidate gene (A), the coordinates for candidate genes (B), and the heat map of LD structure in this region (C). The colored square box in (C) corresponds to each gene with the same color in (B).

We conducted Gene Ontology (GO) enrichment analysis on the significant genes identified by TWASKnockoff using gene expression data from four T2D-related tissues (Figure 7 A, B, C, and D) that showed significantly enriched scores. With gene expression from the sub-cutaneous adipose tissue, TWASKnockoff detected genes associated with cellular metabolic processes, particularly the regulation of the glucose metabolic process (Figure 7 A), which is directly related to the pathogenesis of type 2 diabetes mellitus (T2DM) [36]. Furthermore, genes detected in subcutaneous adipose were enriched in pathways regulating kinase activity, a critical component of cellular functions such as metabolism and insulin signaling [37, 38]. In the pancreas, the identified gene set was significantly enriched in biological pathways associated with the response to osmotic stress and stimulus (Figure 7 C). Osmotic stress, often a result of chronic hyperglycemia, has a profound impact on pancreatic *β*-cell function and survival by increasing reactive oxygen species (ROS) production. This oxidative stress is produced under diabetic conditions and is likely involved in the progression of pancreatic *β*-cell dysfunction found in diabetes [39]. Additionally, the pancreas, particularly the insulin-secreting *β*-cells, is central to responding to endogenous stimuli such as glucose [40] and free fatty acids (FFAs) [41]. Exposure to excessive endogenous stimuli can lead to conditions like Glucotoxicity and Lipotoxicity, leading to insulin resistance, *β*-cell apoptosis, and the progression of T2D [42, 43].

**Figure 7:**
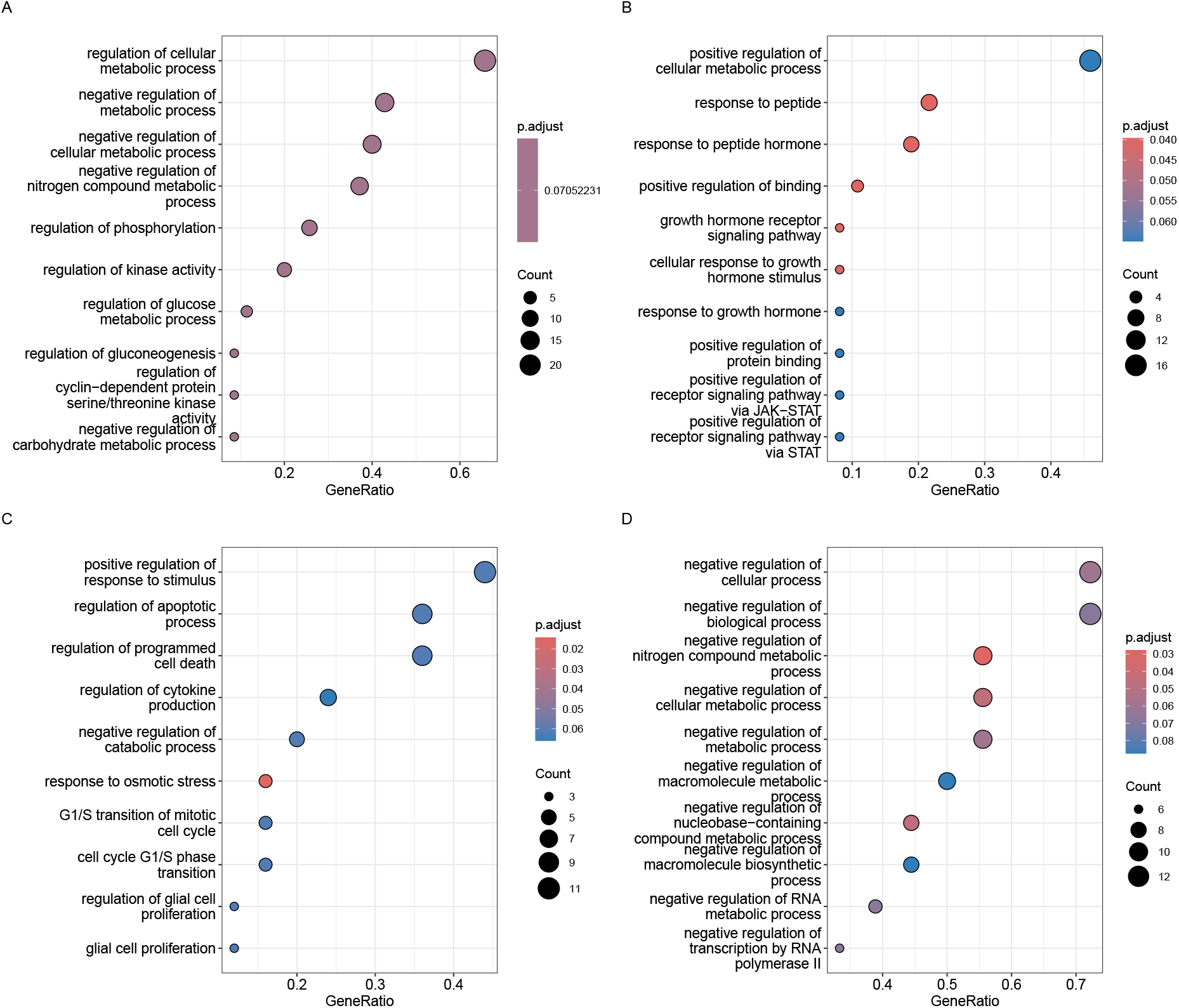
GO analysis of significant genes with FDR *<* 0.1 detected by TWAS-Knockoff for Subcutaneous Adipose (A), Whole Blood (B), Pancreas (C), and Muscle Skeletal (D).

With gene expression data from other T2D-related tissues, TWASKnockoff identified significant genes that are enriched in biologically meaningful pathways. For example, genes detected in whole blood were significantly enriched in pathways related to the response to peptides (Figure 7 B). Specific peptide hormones, such as glucagon-like peptide 1 (GLP-1), play a crucial role in blood sugar regulation by stimulating insulin secretion and suppressing glucagon release [44, 45]. In skeletal muscle cells, TWASKnockoff identified significant genes enriched in pathways regulating the nitrogen compound metabolic process (Figure 7 D). Previous studies have demonstrated the importance of the nitric oxide (NO) pathway in the pathogenesis of diabetes, showing that preserving NO activity could contribute to delaying the onset of insulin resistance and renal dysfunction caused by hyperglycemic stress [46]. Additionally, genes related to the regulation of macromolecule metabolism were also significantly enriched. Previous studies in individuals with and without diabetes have demonstrated the protective effect of an increase in muscle mass on insulin resistance and metabolic syndrome [47, 48, 49]. Skeletal muscle plays a pivotal role in metabolism, serving as a primary target for insulin and the predominant site for insulin-mediated glucose uptake in the postprandial state [50]. Its functions in glucose uptake and energy metabolism, coupled with its importance in exercise and metabolic health, underscore its critical role in metabolic diseases like T2D [51]. These findings demonstrate TWASKnockoff’s ability to identify disease-causing genes with biological functions closely tied to specific tissues, providing insights into tissue-specific pathways underlying T2D.

## 5 Discussion

We present TWASKnockoff, a novel framework for detecting causal gene-tissue pairs by integrating GWAS summary statistics with the gene expression reference panel. TWASKnockoff evaluates the independence of each gene-trait pair, conditional on other genes as well as the direct effects of genetic variants within the same local region. TWASKnockoff first builds gene expression prediction models and then applies the knockoff procedure for variable selection. Additionally, TWASKnockoff enhances the estimation of the correlation matrix for genetic elements using parametric bootstrap samples. Through simulations and applications to the T2D study across ten GTEx tissues, we have demonstrated the benefits of TWASKnockoff in controlling the FDR and improving computational efficiency.

TWASKnockoff offers a flexible, conditional TWAS framework that can be integrated with various statistical methods. For example, the current implementation employs the elastic net to construct gene expression prediction models. However, alternative methods, such as the sum of single-effects (SuSiE) regression model [19], which is an efficient fine-mapping approach, could replace the elastic net to perform the detection of eQTLs and prediction of gene expression levels. Extending TWASKnockoff to incorporate diverse statistical models and comparing their performance in identifying causal gene-tissue pairs is an interesting topic for future research.

Despite its strengths, several challenges remain for further development. First, while TWASKnockoff provides an effective solution for variable selection with a limited number of genes, the use of genetic variants to assist in FDR control can result in overly conservative selections. To address this problem of power loss, we should increase the total number of genes in the TWASKnockoff model. One possible approach is to integrate tissue-specific RNA-seq data to detect causal gene-tissue pairs. This is feasible because TWASKnockoff primarily requires accurate estimation of the correlation matrix for genetic elements, and its computational complexity is largely determined by the number of cis-SNPs in a given region. Consequently, TWASKnockoff can potentially be extended to a highly efficient method for integrating RNA-seq data from multiple tissues and detecting causal gene-tissue pairs. Secondly, as noted in previous studies [52], the inherent randomness of the knockoff method can result in variability in the selected sets of variables across different runs on the same dataset. To mitigate this issue in practical applications, a derandomization approach can be employed, which aggregates selection results across multiple knockoff iterations [53]. Incorporating this strategy into TWASKnockoff could enhance its robustness and reliability.

## 6 Data availability

No data were generated in the present study. We used the UK Biobank genotype data obtained from the UK Biobank resource https://www.ukbiobank.ac.uk/. We used the GTEx data obtained at https://www.ncbi.nlm.nih.gov/projects/gap/cgi-bin/study.cgi?study_id=phs000424.v8.p2. The T2D GWAS summary statistics data are publicly available at http://www.diagram-consortium.org/downloads.html. The 1000 Genomes project data are available at https://www.cog-genomics.org/plink/2.0/resources.

## 7 Code availability

The TWASKnockoff framework is implemented in the R package TWASKnockoff, freely available at https://github.com/zxy0912/TWASKnockoff.

## 8 Acknowledgment

This work was supported in part by NIH grants R01 GM134005 and U24 HG012108. We thank the participants of the UK Biobank and the GTEx (v8) project.

## References

[1] Peter M Visscher, Matthew A Brown, Mark I McCarthy, and Jian Yang. Five years of gwas discovery. The American Journal of Human Genetics, 90(1):7–24, 2012.

[2] Michael D Gallagher and Alice S Chen-Plotkin. The post-gwas era: from association to function. The American Journal of Human Genetics, 102(5):717–730, 2018.

[3] Eric R Gamazon, Heather E Wheeler, Kaanan P Shah, Sahar V Mozaffari, Keston Aquino-Michaels, Robert J Carroll, Anne E Eyler, Joshua C Denny, GTEx Consortium, Dan L Nicolae, et al. A gene-based association method for mapping traits using reference transcriptome data. Nature genetics, 47(9):1091–1098, 2015.

[4] Alexander Gusev, Arthur Ko, Huwenbo Shi, Gaurav Bhatia, Wonil Chung, Brenda WJH Penninx, Rick Jansen, Eco JC De Geus, Dorret I Boomsma, Fred A Wright, et al. Integrative approaches for large-scale transcriptome-wide association studies. Nature genetics, 48(3):245–252, 2016.

[5] Michael Wainberg, Nasa Sinnott-Armstrong, Nicholas Mancuso, Alvaro N Barbeira, David A Knowles, David Golan, Raili Ermel, Arno Ruusalepp, Thomas Quertermous, Ke Hao, et al. Opportunities and challenges for transcriptome-wide association studies. Nature genetics, 51(4):592–599, 2019.

[6] Tiffany Amariuta, Katherine Siewert-Rocks, and Alkes L Price. Modeling tissue co-regulation estimates tissue-specific contributions to disease. Nature genetics, 55(9):1503–1511, 2023.

[7] Nicholas Mancuso, Malika K Freund, Ruth Johnson, Huwenbo Shi, Gleb Kichaev, Alexander Gusev, and Bogdan Pasaniuc. Probabilistic fine-mapping of transcriptome-wide association studies. Nature genetics, 51(4):675–682, 2019.

[8] Chong Wu and Wei Pan. A powerful fine-mapping method for transcriptome-wide association studies. Human genetics, 139(2):199–213, 2020.

[9] Douglas W Yao, Luke J O’connor, Alkes L Price, and Alexander Gusev. Quantifying genetic effects on disease mediated by assayed gene expression levels. Nature genetics, 52(6):626–633, 2020.

[10] Siming Zhao, Wesley Crouse, Sheng Qian, Kaixuan Luo, Matthew Stephens, and Xin He. Adjusting for genetic confounders in transcriptome-wide association studies leads to reliable detection of causal genes. bioRxiv, pages 2022–09, 2022.

[11] Lu Liu, Ran Yan, Ping Guo, Jiadong Ji, Weiming Gong, Fuzhong Xue, Zhongshang Yuan, and Xiang Zhou. Conditional transcriptome-wide association study for fine-mapping candidate causal genes. Nature Genetics, pages 1–9, 2024.

[12] Rina Foygel Barber and Emmanuel J Candès. Controlling the false discovery rate via knockoffs. The Annals of statistics, pages 2055–2085, 2015.

[13] Emmanuel Candes, Yingying Fan, Lucas Janson, and Jinchi Lv. Panning for gold:’model-x’knockoffs for high dimensional controlled variable selection. Journal of the Royal Statistical Society Series B: Statistical Methodology, 80(3):551–577, 2018.

[14] Matteo Sesia, Eugene Katsevich, Stephen Bates, Emmanuel Candès, and Chiara Sabatti. Multi-resolution localization of causal variants across the genome. Nature communications, 11(1):1093, 2020.

[15] Matteo Sesia, Stephen Bates, Emmanuel Candès, Jonathan Marchini, and Chiara Sabatti. False discovery rate control in genome-wide association studies with population structure. Proceedings of the National Academy of Sciences, 118(40):e2105841118, 2021.

[16] Anqi Wang, Peixin Tian, and Yan Dora Zhang. Twas-gkf: a novel method for causal gene identification in transcriptome-wide association studies with knockoff inference. Bioinformatics, 40(8):btae502, 2024.

[17] Zihuai He, Linxi Liu, Michael E Belloy, Yann Le Guen, Aaron Sossin, Xiaoxia Liu, Xinran Qi, Shiyang Ma, Prashnna K Gyawali, Tony Wyss-Coray, et al. Ghostknockoff inference empowers identification of putative causal variants in genome-wide association studies. Nature communications, 13(1):7209, 2022.

[18] Zhaomeng Chen, Zihuai He, Benjamin B Chu, Jiaqi Gu, Tim Morrison, Chiara Sabatti, and Emmanuel Candès. Controlled variable selection from summary statistics only? a solution via ghostknockoffs and penalized regression. ArXiv, 2024.

[19] Gao Wang, Abhishek Sarkar, Peter Carbonetto, and Matthew Stephens. A simple new approach to variable selection in regression, with application to genetic fine mapping. Journal of the Royal Statistical Society Series B: Statistical Methodology, 82(5):1273–1300, 2020.

[20] Genomes Project Consortium, A Auton, LD Brooks, RM Durbin, EP Garrison, and HM Kang. A global reference for human genetic variation. Nature, 526(7571):68–74, 2015.

[21] GTEx Consortium. The gtex consortium atlas of genetic regulatory effects across human tissues. Science, 369(6509):1318–1330, 2020.

[22] Ken Suzuki, Konstantinos Hatzikotoulas, Lorraine Southam, Henry J Taylor, Xianyong Yin, Kim M Lorenz, Ravi Mandla, Alicia Huerta-Chagoya, Giorgio EM Melloni, Stavroula Kanoni, et al. Genetic drivers of heterogeneity in type 2 diabetes pathophysiology. Nature, 627(8003):347–357, 2024.

[23] Gil Stelzer, Naomi Rosen, Inbar Plaschkes, Shahar Zimmerman, Michal Twik, Simon Fishilevich, Tsippi Iny Stein, Ron Nudel, Iris Lieder, Yaron Mazor, et al. The genecards suite: from gene data mining to disease genome sequence analyses. Current protocols in bioinformatics, 54(1):1–30, 2016.

[24] Struan FA Grant, Gudmar Thorleifsson, Inga Reynisdottir, Rafn Benediktsson, Andrei Manolescu, Jesus Sainz, Agnar Helgason, Hreinn Stefansson, Valur Emilsson, Anna Helgadottir, et al. Variant of transcription factor 7-like 2 (tcf7l2) gene confers risk of type 2 diabetes. Nature genetics, 38(3):320–323, 2006.

[25] Laura del Bosque-Plata, Eduardo Martínez-Martínez, Miguel Ángel Espinoza-Camacho, and Claudia Gragnoli. The role of tcf7l2 in type 2 diabetes. Diabetes, 70(6):1220–1228, 2021.

[26] Valeriya Lyssenko, Roberto Lupi, Piero Marchetti, Silvia Del Guerra, Marju Orho-Melander, Peter Almgren, Marketa Sjögren, Charlotte Ling, Karl-Fredrik Eriksson, Rita Mancarella, et al. Mechanisms by which common variants in the tcf7l2 gene increase risk of type 2 diabetes. The Journal of clinical investigation, 117(8):2155–2163, 2007.

[27] Muhammad Sajid Hamid Akash, Kanwal Rehman, and Aamira Liaqat. Tumor necrosis factor-alpha: role in development of insulin resistance and pathogenesis of type 2 diabetes mellitus. Journal of cellular biochemistry, 119(1):105–110, 2018.

[28] Hana Alzamil. Elevated serum tnf-α is related to obesity in type 2 diabetes mellitus and is associated with glycemic control and insulin resistance. Journal of obesity, 2020(1):5076858, 2020.

[29] Valgerdur Steinthorsdottir, Gudmar Thorleifsson, Patrick Sulem, Hannes Helgason, Niels Grarup, Asgeir Sigurdsson, Hafdis T Helgadottir, Hrefna Johannsdottir, Olafur T Magnusson, Sigurjon A Gudjonsson, et al. Identification of low-frequency and rare sequence variants associated with elevated or reduced risk of type 2 diabetes. Nature genetics, 46(3):294–298, 2014.

[30] Senta Georgia, Anil Bhushan, et al. β cell replication is the primary mechanism for maintaining postnatal β cell mass. The Journal of clinical investigation, 114(7):963–968, 2004.

[31] Jake A Kushner, Maria A Ciemerych, Ewa Sicinska, Lynn M Wartschow, Monica Teta, Simon Y Long, Piotr Sicinski, and Morris F White. Cyclins d2 and d1 are essential for postnatal pancreatic β-cell growth. Molecular and cellular biology, 25(9):3752–3762, 2005.

[32] Junguo Cao, Weijia Yan, Xiujian Ma, Haiyan Huang, and Hong Yan. Insulin-like growth factor 2 mrna-binding protein 2—a potential link between type 2 diabetes mellitus and cancer. The journal of clinical endocrinology & metabolism, 106(10):2807–2818, 2021.

[33] Jinyan Wang, Lijuan Chen, and Ping Qiang. The role of igf2bp2, an m6a reader gene, in human metabolic diseases and cancers. Cancer cell international, 21(1):99, 2021.

[34] Tianwei Gu, Eva Horová, Anna Möllsten, Norhashimah Abu Seman, Henrik Falhammar, Martin Prázny, Kerstin Brismar, and Harvest F Gu. Igf2bp2 and igf2 genetic effects in diabetes and diabetic nephropathy. Journal of diabetes and its complications, 26(5):393–398, 2012.

[35] Constanze M Hammerle, Ionel Sandovici, Gemma V Brierley, Nicola M Smith, Warren E Zimmer, Ilona Zvetkova, Haydn M Prosser, Yoichi Sekita, Brian YH Lam, Marcella Ma, et al. Mesenchyme-derived igf2 is a major paracrine regulator of pancreatic growth and function. PLoS Genetics, 16(10):e1009069, 2020.

[36] Ralph A DeFronzo. Pathogenesis of type 2 diabetes mellitus. Medical clinics, 88(4):787–835, 2004.

[37] Simon M Schultze, Brian A Hemmings, Markus Niessen, and Oliver Tschopp. Pi3k/akt, mapk and ampk signalling: protein kinases in glucose homeostasis. Expert reviews in molecular medicine, 14:e1, 2012.

[38] Sanyogita Chauhan, Aakash Partap Singh, Avtar Chand Rana, Sunil Kumar, Ravi Kumar, Jitender Singh, Ashok Jangra, and Dinesh Kumar. Natural activators of ampk signaling: potential role in the management of type-2 diabetes. Journal of Diabetes & Metabolic Disorders, 22(1):47–59, 2023.

[39] Natsuki Eguchi, Nosratola D Vaziri, Donald C Dafoe, and Hirohito Ichii. The role of oxidative stress in pancreatic β cell dysfunction in diabetes. International journal of molecular sciences, 22(4):1509, 2021.

[40] R Paul Robertson, Jamie Harmon, Phuong Oanh Tran, Yoshito Tanaka, and Hiroki Takahashi. Glucose toxicity in β-cells: type 2 diabetes, good radicals gone bad, and the glutathione connection. Diabetes, 52(3):581–587, 2003.

[41] Yoon S Oh, Gong D Bae, Dong J Baek, Eun-Young Park, and Hee-Sook Jun. Fatty acid-induced lipotoxicity in pancreatic beta-cells during development of type 2 diabetes. Frontiers in endocrinology, 9:384, 2018.

[42] Ji-Won Kim and Kun-Ho Yoon. Glucolipotoxicity in pancreatic β-cells. Diabetes & metabolism journal, 35(5):444–450, 2011.

[43] Dilek Yazıcı and Havva Sezer. Insulin resistance, obesity and lipotoxicity. Obesity and lipotoxicity, pages 277–304, 2017.

[44] Deborah Hinnen. Glucagon-like peptide 1 receptor agonists for type 2 diabetes. Diabetes spectrum, 30(3):202–210, 2017.

[45] Michael A Nauck and Timo D Müller. Incretin hormones and type 2 diabetes. Diabetologia, 66(10):1780–1795, 2023.

[46] Taylor Claybaugh, Sarah Decker, Kelly McCall, Yuriy Slyvka, Jerrod Steimle, Aaron Wood, Megan Schaefer, Jean Thuma, and Sharon Inman. L-arginine supplementation in type ii diabetic rats preserves renal function and improves insulin sensitivity by altering the nitric oxide pathway. International journal of endocrinology, 2014(1):171546, 2014.

[47] Naomi Brooks, Jennifer E Layne, Patricia L Gordon, Ronenn Roubenoff, Miriam E Nelson, and Carmen Castaneda-Sceppa. Strength training improves muscle quality and insulin sensitivity in hispanic older adults with type 2 diabetes. International journal of medical sciences, 4(1):19, 2007.

[48] Lee D Katz, Morton G Glickman, Stanle Rapoport, Eleuterio Ferrannini, and Ralph A DeFronzo. Splanchnic and peripheral disposal of oral glucose in man. Diabetes, 32(7):675–679, 1983.

[49] Kyuwoong Kim and Sang Min Park. Association of muscle mass and fat mass with insulin resistance and the prevalence of metabolic syndrome in korean adults: a cross-sectional study. Scientific reports, 8(1):2703, 2018.

[50] Ralph A DeFronzo and Devjit Tripathy. Skeletal muscle insulin resistance is the primary defect in type 2 diabetes. Diabetes care, 32(Suppl 2):S157, 2009.

[51] Karla E Merz and Debbie C Thurmond. Role of skeletal muscle in insulin resistance and glucose uptake. Comprehensive Physiology, 10(3):785–809, 2011.

[52] Zhimei Ren and Rina Foygel Barber. Derandomised knockoffs: leveraging e-values for false discovery rate control. Journal of the Royal Statistical Society Series B: Statistical Methodology, 86(1):122–154, 2024.

[53] Zhimei Ren, Yuting Wei, and Emmanuel Candès. Derandomizing knockoffs. Journal of the American Statistical Association, 118(542):948–958, 2023.

