## Supplementary materials for "Knockoff procedure improves causal gene identifications in conditional transcriptome-wide association studies"

Xiangyu Zhang[1], Lijun Wang[1], Jia Zhao[1], Hongyu Zhao[1*]

**1** Department of Biostatistics, School of Public Health, Yale University, New Haven, Connecticut, United States of America

#### S1 FDR control via TWASKnockoff

Suppose $Z = [z_{\hat{G}}^T, z_X^T]^T$ contains the z-scores for all genetic elements. To identify variables relevant to the trait of interest, we compute feature statistics $T_j$ for each covariate as follows:

$$T_j = T_j([Z, \tilde{Z}], y),$$

where $\tilde{Z}$ is the knockoff copy of the z-scores.

For a fixed threshold $t > 0$, we have

$$\#\{j : T_j \leq -t\} \geq \#\{null\ j : T_j \leq -t\} \geq \#\{null\ j : T_j \geq t\}.$$

We select variables such that their feature statistics are sufficiently large. The FDR

$$\mathrm{FDR}(t) = \frac{\#\{null\ j : T_j \geq t\}}{\#\{j : T_j \geq t\}}$$

could be controlled by:

$$\widehat{FDR}(t) = \frac{\#\{j : T_j \leq -t\}}{\#\{j : T_j \geq t\}}.$$

Next, we aim to control the FDR within the subset of candidate genes. Let $A$ denote the set of indices for candidate genes and $B$ denote the set of indices for candidate SNPs. The gene-level FDR is given by:

$$FDR_{gene}(t) = \frac{\#\{null\ j \in A : T_j \geq t\}}{\#\{j \in A : T_j \geq t\}}.$$

To simplify, consider a case where there are no non-mediated causal SNPs, in other words, all variables from set $B$ are nulls. With the properties of feature statistics:

$$\#\{null\ j \in B : T_j \geq t\} = \#\{j \in B : T_j \geq t\}.$$

Thus, since it always holds that:

$$\#\{null\ j \in A : T_j \geq t\} \leq \#\{j \in A : T_j \geq t\},$$

we have the following for the gene-level FDR:

$$
\begin{aligned}
FDR_{gene}(t) =& \frac{\#\{null\ j \in A : T_j \geq t\}}{\#\{j \in A : T_j \geq t\}} \\
\leq& \frac{\#\{null\ j \in A : T_j \geq t\} + \#\{null\ j \in B : T_j \geq t\}}{\#\{j \in A : T_j \geq t\} + \#\{j \in B : T_j \geq t\}} \\
=& \frac{\#\{null\ j : T_j \geq t\}}{\#\{j : T_j \geq t\}} \\
\leq& \frac{\#\{j : T_j \leq -t\}}{\#\{j : T_j \geq t\}} = \widehat{FDR}(t).
\end{aligned} \tag{1}
$$

The first inequality always holds if the following property holds:

$$\frac{\#\{null\ j \in A : T_j \geq t\}}{\#\{j \in A : T_j \geq t\}} \leq \frac{\#\{null\ j \in B : T_j \geq t\}}{\#\{j \in B : T_j \geq t\}}.$$

This suggests that we can control the FDR for candidate genes at a level below the estimated FDR for all genetic elements. Since the probability of a candidate SNP being causal for the trait of interest is generally much lower than the probability of a candidate gene being causal, this condition is typically satisfied in most cases.

To verify this condition in real data application, we can evaluate whether the following condition is satisfied:

$$\frac{\#\{j \in A : T_j \leq -t\}}{\#\{j \in A : T_j \geq t\}} \leq \frac{\#\{j \in B : T_j \leq -t\}}{\#\{j \in B : T_j \geq t\}},$$

which leads to the same inequality:

$$
\begin{aligned}
FDR_{gene}(t) =& \frac{\#\{null\ j \in A : T_j \geq t\}}{\#\{j \in A : T_j \geq t\}} \\
\leq& \frac{\#\{j \in A : T_j \leq -t\}}{\#\{j \in A : T_j \geq t\}} \\
\leq& \frac{\#\{j \in A : T_j \leq -t\} + \#\{j \in B : T_j \leq -t\}}{\#\{j \in A : T_j \geq t\} + \#\{j \in B : T_j \geq t\}} \\
=& \frac{\#\{j : T_j \leq -t\}}{\#\{j : T_j \geq t\}} = \widehat{FDR}(t).
\end{aligned} \tag{2}
$$

This derivation confirms that the FDR for candidate genes can be effectively bounded by the estimated FDR for all genetic elements.

#### S2 Improved Estimation of Correlation Matrix

When calculating the covariance matrix of $[\hat{\mathbf{G}}\ \mathbf{X}_t]$, where $\hat{\mathbf{G}} = \mathbf{X}_t\hat{\boldsymbol{\gamma}}$, we proposed incorporating the randomness of $\hat{\boldsymbol{\gamma}}$. Denote the naive estimates without taking the randomness of $\hat{\boldsymbol{\gamma}}$ into account as:

$$
\widehat{\text{Cov}}_{\text{Naive}}(\hat{\mathbf{G}}) = \hat{\boldsymbol{\gamma}}^T\hat{\mathbf{R}}\boldsymbol{\gamma}\,, \qquad \widehat{\text{Cov}}_{\text{Naive}}(\hat{\mathbf{G}}, \mathbf{X}_t) = \hat{\boldsymbol{\gamma}}^T\hat{\mathbf{R}}\,.
$$

We approximate the underlying true expectation by $N$ Monte Carlo samples:

$$
\widehat{\text{Cov}}_{\text{MC}}(\hat{\mathbf{G}}) = \frac{1}{N}\sum_{i=1}^{N}\hat{\boldsymbol{\gamma}}^{(i)T}\hat{\mathbf{R}}\hat{\boldsymbol{\gamma}}^{(i)}\,, \qquad \widehat{\text{Cov}}_{\text{MC}}(\hat{\mathbf{G}}, \mathbf{X}_t) = \frac{1}{N}\sum_{i=1}^{N}\hat{\boldsymbol{\gamma}}^{(i)T}\hat{\mathbf{R}}\,,
$$

where

$$
\hat{\boldsymbol{\gamma}}^{(i)} = \hat{f}(\mathbf{X}_e, \mathbf{X}_e\boldsymbol{\gamma}_0 + \mathbf{e}^{(i)})\,,
$$

and $\mathbf{e}^{(i)}$ is resampling from the true data-generating model.

In practice, we do not know the data-generating model, instead, we take the parametric bootstrap procedure to incorporate the randomness of $\hat{\boldsymbol{\gamma}}$:

$$\widehat{\mathrm{Cov}}_{\mathrm{Bootstrap}}(\hat{\mathbf{G}}) = \frac{1}{B}\sum_{b=1}^{B} \hat{\boldsymbol{\gamma}}_\star^{(b)T}\hat{\mathbf{R}}\hat{\boldsymbol{\gamma}}_\star^{(b)}\,, \qquad \widehat{\mathrm{Cov}}_{\mathrm{Bootstrap}}(\hat{\mathbf{G}}, \mathbf{X}_t) = \frac{1}{B}\sum_{b=1}^{B} \hat{\boldsymbol{\gamma}}_\star^{(b)T}\hat{\mathbf{R}}\,,$$

where

$$\hat{\gamma}_\star^{(b)} = \hat{f}(\mathbf{X}_e, \mathbf{X}_e\hat{\boldsymbol{\gamma}} + \hat{\mathbf{e}}_\star^{(b)})\,,$$

and $\mathbf{e}_\star^{(b)}$ is the $b$-th bootstrapped error term.

Note that a correlation matrix is obtained by standardizing the covariance matrix, and our procedure will finally handle the correlation matrix, so we directly check the improvement of the estimates of correlation matrices. We define the gap as the Frobenius norm of the difference of two correlation matrices:

$$\begin{aligned}
\mathrm{Gap}_{\text{Gene-Gene}}(\mathrm{MC}, \mathrm{Naive}) &= \|\widehat{\mathrm{Cor}}_{\mathrm{MC}}(\hat{\mathbf{G}}) - \widehat{\mathrm{Cor}}_{\mathrm{Naive}}(\hat{\mathbf{G}})\|_F\,,\\
\mathrm{Gap}_{\text{Gene-Gene}}(\mathrm{MC}, \mathrm{Bootstrap}) &= \|\widehat{\mathrm{Cor}}_{\mathrm{MC}}(\hat{\mathbf{G}}) - \widehat{\mathrm{Cor}}_{\mathrm{Bootstrap}}(\hat{\mathbf{G}})\|_F\,,\\
\mathrm{Gap}_{\text{Gene-SNP}}(\mathrm{MC}, \mathrm{Naive}) &= \|\widehat{\mathrm{Cor}}_{\mathrm{MC}}(\hat{\mathbf{G}}, \mathbf{X}_t) - \widehat{\mathrm{Cor}}_{\mathrm{Naive}}(\hat{\mathbf{G}}, \mathbf{X}_t)\|_F\,,\\
\mathrm{Gap}_{\text{Gene-SNP}}(\mathrm{MC}, \mathrm{Bootstrap}) &= \|\widehat{\mathrm{Cor}}_{\mathrm{MC}}(\hat{\mathbf{G}}, \mathbf{X}_t) - \widehat{\mathrm{Cor}}_{\mathrm{Bootstrap}}(\hat{\mathbf{G}}, \mathbf{X}_t)\|_F\,.
\end{aligned}$$

Figures 1a and 1b illustrate the gap between the Monte Carlo approximation and the bootstrap estimate as a function of the number of bootstrap samples $B$, for the gene-gene correlation and the gene-SNP correlation, respectively. In both cases, the gap between the Monte Carlo approximation and the naive estimate remains constant. Figures 1c and 1d show the gaps in 100 repetitive experiments 100 times, each conducted with $N = 100$ Monte Carlo samples and $B = 10$ bootstrap samples. The results demonstrate that incorporating the randomness of $\hat{\boldsymbol{\gamma}}$ can achieve a more accurate and more stable estimate for both gene-gene and gene-SNP correlations. While increasing the number of bootstrap samples could further enhance accuracy and stability, we chose $B = 10$ in Figures 1c and 1d, as well as in Section **??**, to maintain a balance between computational efficiency and performance.

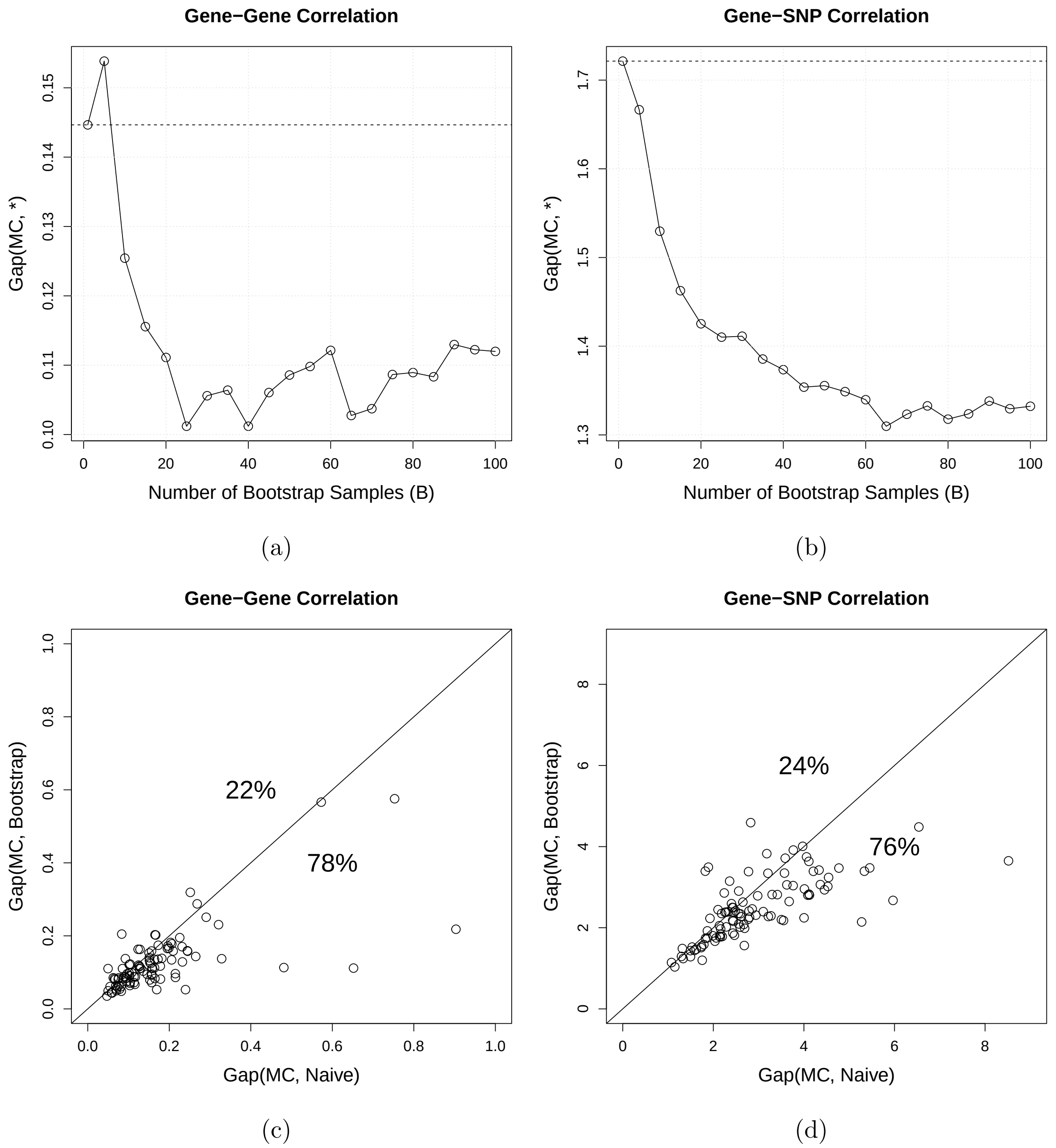


Figure 1: ***(Top)*** The gap between the Monte Carlo approximation ($N = 100$) and the bootstrap estimate versus the number of bootstrap samples $B$ for the gene-gene correlation ***(a)*** and the gene-SNP correlation ***(b)***, respectively. The dashed horizontal line indicates the constant gap between the Monte Carlo approximation and the naive approach. ***(Bottom)*** Gaps of the naive approach and the bootstrap ($B = 10$) estimate compared to the Monte Carlo approximation ($N = 100$) for the gene-gene correlation ***(c)*** and the gene-SNP correlation ***(d)***, respectively. The percentage indicates the proportion in which one is larger than the other among 100 experiments.

### S3 Data processing

#### S3.1 UKBB genotype

We extracted overlapping SNPs from UKBB, 1000 Genomes Project Phase 3 (1KG Phase 3), and HapMap3 [1] datasets to perform simulations. For this study, we selected genetically unrelated participants of White British ancestry ($n = 276,731$). We applied quality control to the UKBB data with European ancestry using PLINK [2]. To be more specific, we excluded samples with more than 10% missing genotypes and markers that failed the Hardy-Weinberg test, we included only SNPs with a genotyping rate of at least 95% and a minor allele frequency (MAF) of at least 1% (–geno 0.05 –hwe 1e-10 –mind 0.1 –maf 0.01). After these quality control steps, the final genotype dataset included 276,731 unrelated European individuals genotyped at 1,126,249 SNPs.

#### S3.2 1KG reference panel

We downloaded 1000 Genomes (1KG) phase 3 reference panels for GRCH37/hg19 (used as the external reference panel for simulations) and GRCH38/hg38 (used as the external for real data analysis) from `https://www.cog-genomics.org/plink/2.0/resources`. Using the superpopulation information provided by the 1000 Genomes Project, we selected European samples and excluded all duplicated and ambiguous SNPs. Quality control was applied to the 1KG reference panel for individuals of European ancestry using PLINK [2]. To be more specific, we excluded samples with more than 10% missing genotypes and markers that failed the Hardy-Weinberg test, we only included SNPs with at least 95% genotyping rate with minor allele frequency (MAF) no less than 1% (–geno 0.05 –hwe 1e-10 –mind 0.1 –maf 0.01). For build 37, the final genotype data includes 503 non-overlapping Europeans genotyped at 7,222,123 SNPs. For build 38, the final genotype data includes 633 non-overlapping Europeans genotyped at 7,412,727 SNPs.

##### S3.3 T2D GWAS summary statistics

We used summary statistics for T2D obtained from a recent large meta-analysis of T2D GWAS data comprising more than 2.5 million individuals of diverse ancestry [3]. The data can be downloaded from `http://www.diagram-consortium.org/downloads.html`. We focused on the European ancestry group (60.3% of the effective sample size) and used the liftOver tool (`https://genome-store.ucsc.edu`) to convert SNP coordinates to build GRCh38/hg38. The final dataset included summary statistics at 19,296,716 SNPs, with a median of sample size 532,776.

##### S3.4 GTEx RNA-seq data

We performed quality control on the genotype data from the GTEx (v8) project using PLINK [2]. Samples with more than 10% missing genotypes and markers failing the Hardy-Weinberg equilibrium test were excluded. SNPs were retained if they had a genotyping rate of at least 95% and a minor allele frequency (MAF) of at least 1% (–geno 0.05, –hwe 1e-6, –mind 0.1, –maf 0.01). The final dataset comprised 837 individuals and 10,520,159 SNPs. Expression data from GTEx (v8), encompassing 948 individuals, 56,200 genes, and 54 tissues, were adjusted for covariates including sex, age, the first five genotype principal components (PCs), and the top five probabilistic estimation of expression residuals (PEER factors), to account for potential confounding effects.

##### S3.5 GTEx genotype data

Using the meta-data information provided by the GTEx Project, we selected European samples and excluded all duplicated and ambiguous SNPs. Quality control was applied to the GTEx genotype data using PLINK [2]. To be more specific, we excluded samples with more than 10% missing genotypes and markers that failed the Hardy-Weinberg test, we only included SNPs with at least 95% genotyping rate with minor allele frequency (MAF) no less than 1% (–geno 0.05 –hwe 1e-10 –mind 0.1 –maf 0.01). The final genotype data includes 715 non-overlapping Europeans genotyped at 6,937,325 SNPs.

#### S4 Details for methods compared

**GIFT**: We ran GIFT in all simulations by calling functions from GIFT v2.0 (`https://github.com/yuanzhongshang/GIFT/tree/v2.0`). For simulations with the individual level data, we called the function `GIFT_individual` function. For simulations with summary statistics and an external reference panel, as well as the real data analysis, we called the `GIFT_summary` function. To ensure the convergence of the algorithm, the maximum iteration was fixed at 100 in simulations and 1,000 in real data analysis, while the tolerance of the absolute value of the difference between $n$ th and $(n+1)$ th log-likelihood was fixed at $1e-3$ in simulations and $1e-4$ in real data analysis.

**GhostKnockoff**: We applied GhostKnockoff v0.1.0 (`https://github.com/cran/GhostKnockoff/`, `https://github.com/biona001/ghostknockoff-gwas-reproducibility/tree/main/chen_et_al`) to perform variable selection in TWASKnockoff. We called the function `GhostKnockoff.prelim` and `GhostKnockoff.fit` to generate knockoff copies and set the number of knockoff copies per variant at $M = 1$. We then called the function `GhostKnockoff.filter` to perform variable selection. We tested various feature statistics within TWASKnockoff, including lasso coefficients (`lasso`), lasso coefficients with approximated $\lambda$ (`lasso.approx.lambda`), marginal empirical correlations (`marginal`), the element-wise square of z-scores (`squared.zscore`), and posterior inclusion probability produced by SuSiE (`susie`).

#### S5 Supplementary plots

#### References

[1] Richard A Gibbs, John W Belmont, Paul Hardenbol, Thomas D Willis, Fuli L Yu, HM Yang, Lan-Yang Ch'ang, Wei Huang, Bin Liu, Yan Shen, et al. The international hapmap project. 2003.

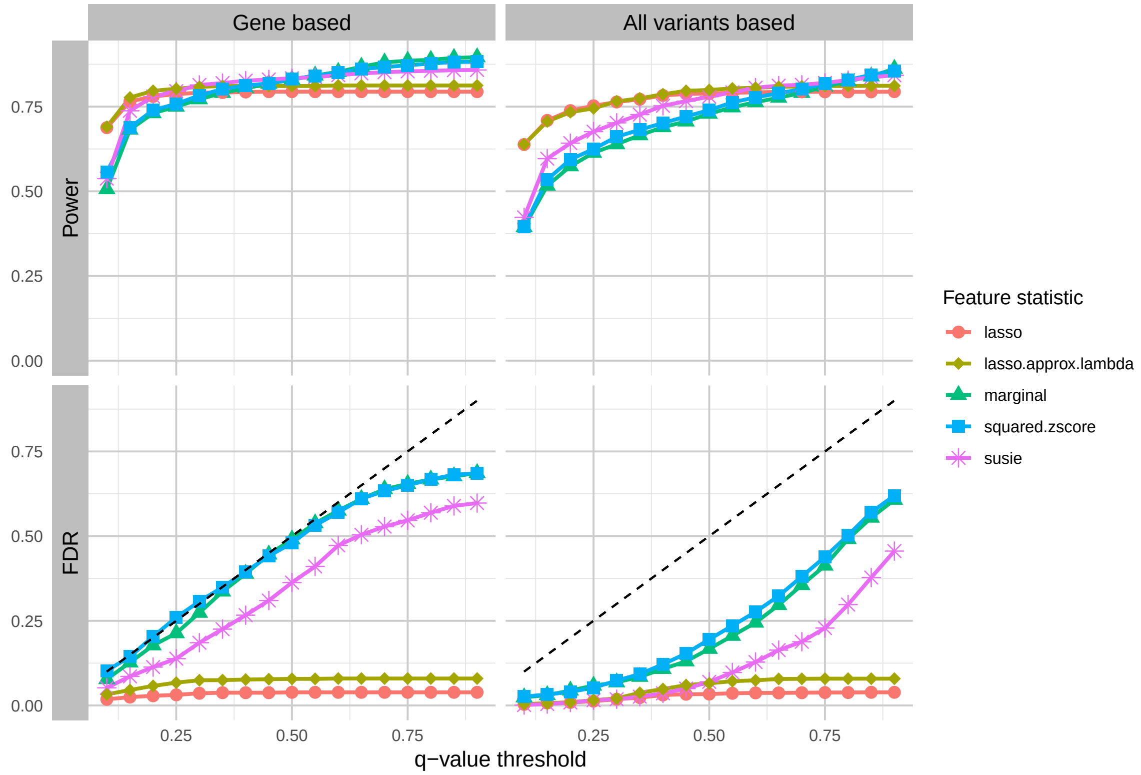


Figure 2: **Comparison between two FDR control strategies.** The FDR of genes was controlled within genes only (left) and all genetic elements (right) in TWASKnockoff, with the empirical estimation of the correlation matrix. In this simulation, all the heritability was explained by causal genes within the risk region, and we considered five feature statistics in the GhostKnockoff framework.

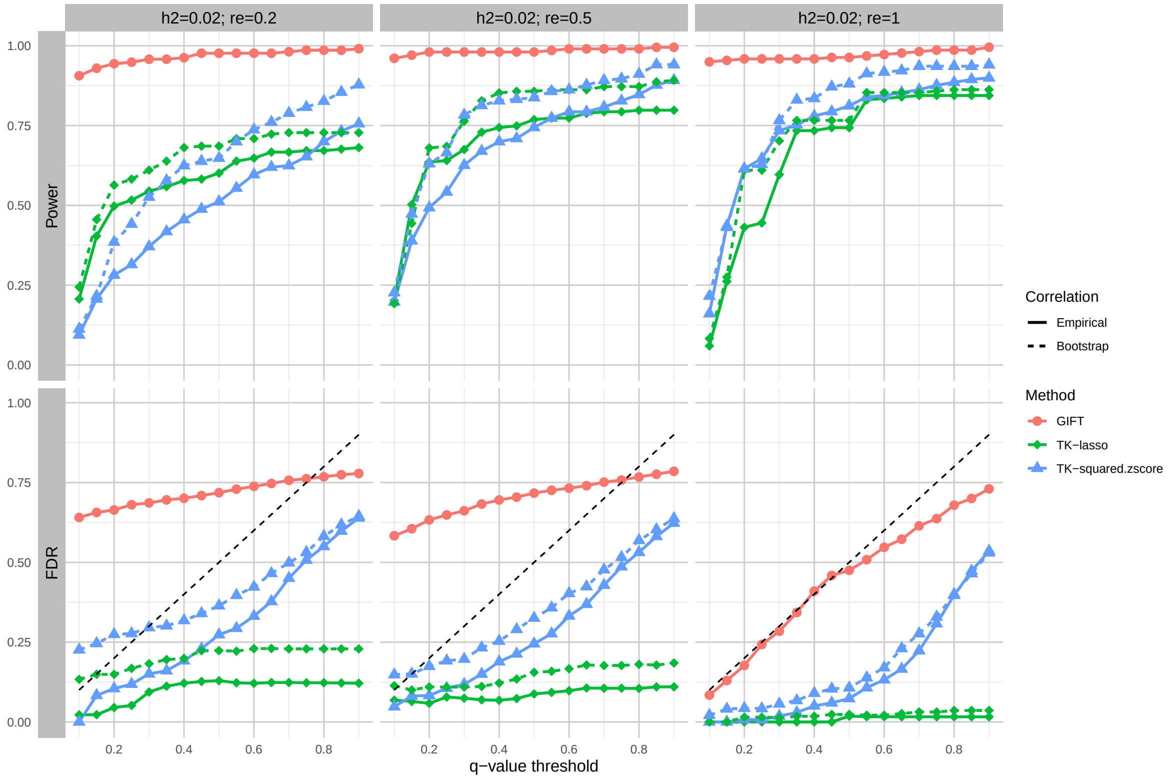


Figure 3: **Comparison of TWASKnockoff and GIFT in simulated data with the in-sample LD matrix when** $h^2 = 0.02$**.** We assess the performance of GIFT and TWAS-Knockoff with two feature statistics: lasso coefficients (TK-lasso) and squared z-scores (TK-squared.zscore). For each feature statistic, we consider two estimation methods (empirical estimation and bootstrap samples) of the correlation matrix of genetic elements. The $q$-value threshold is selected from 0.1 to 0.9, with the black dashed line indicating the theoretical $q$-value level. For GIFT, we applied the Benjamini-Hochberg (BH) correction to perform FDR control.

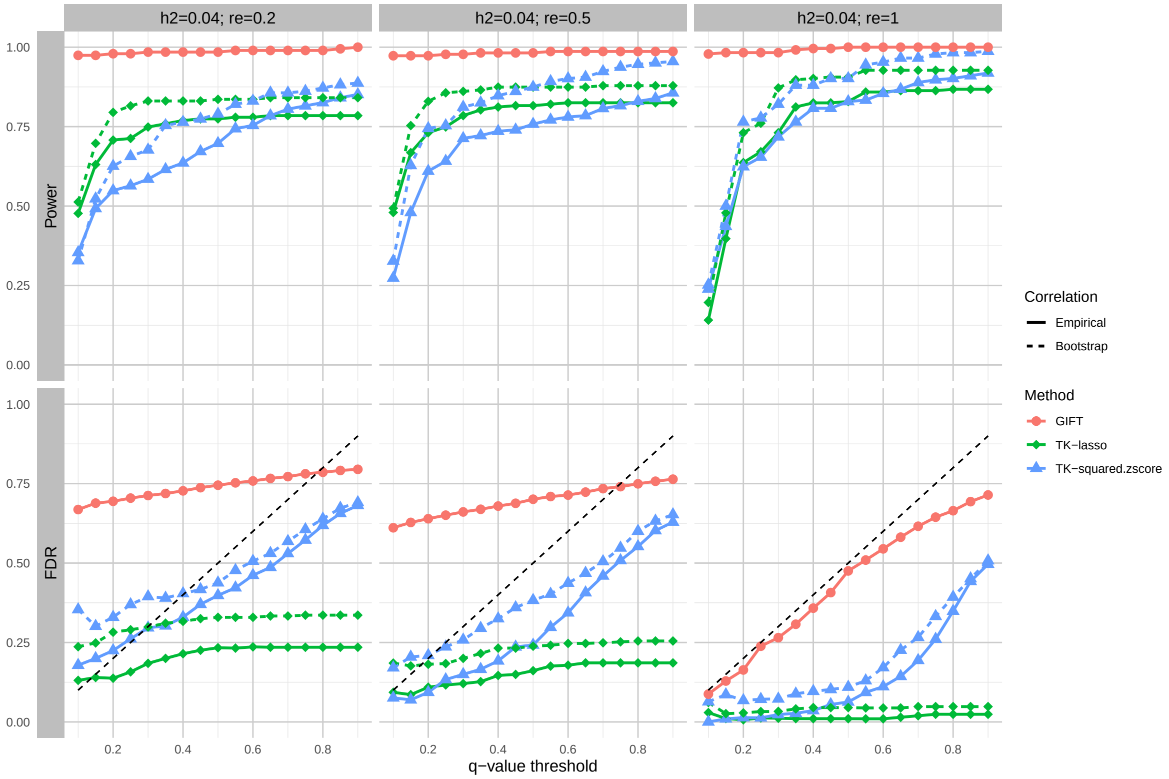


Figure 4: **Comparison of TWASKnockoff and GIFT in simulated data with the in-sample LD matrix when $h^2 = 0.04$.** We assess the performance of GIFT and TWAS-Knockoff with two feature statistics: lasso coefficients (TK-lasso) and squared z-scores (TK-squared.zscore). For each feature statistic, we consider two estimation methods (empirical estimation and bootstrap samples) of the correlation matrix of genetic elements. The $q$-value threshold is selected from 0.1 to 0.9, with the black dashed line indicating the theoretical $q$-value level. For GIFT, we applied the Benjamini-Hochberg (BH) correction to perform FDR control.

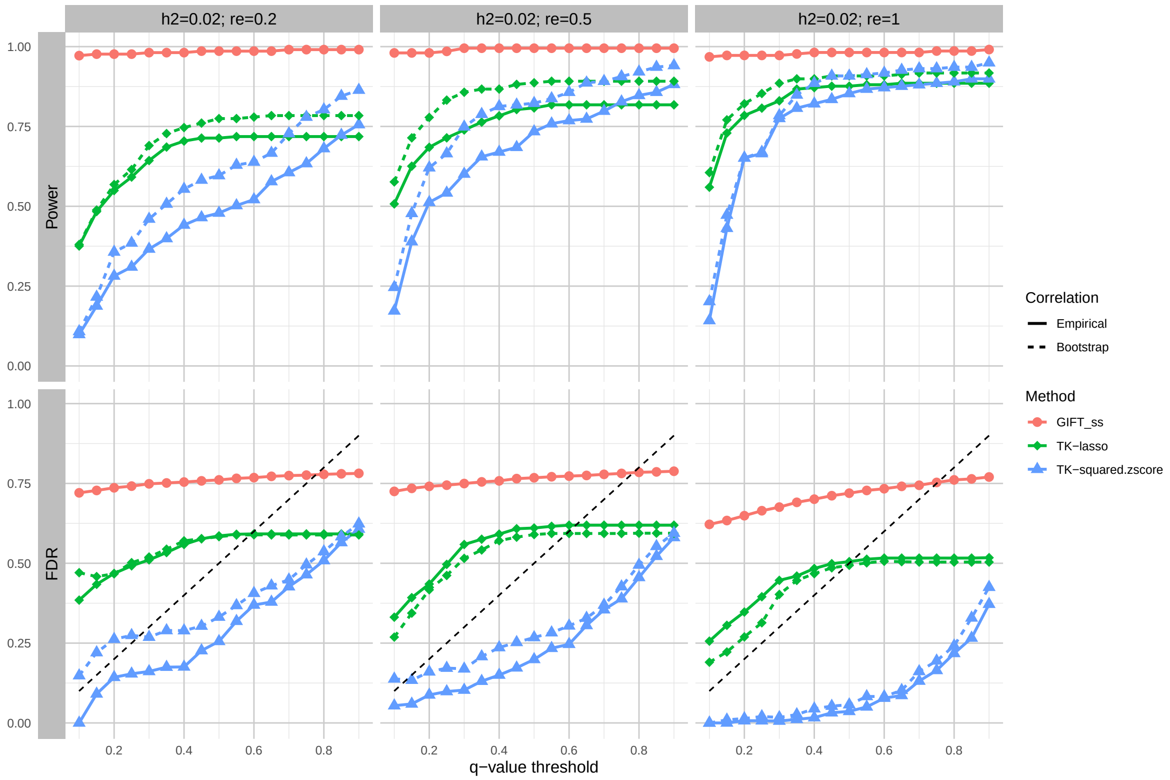


Figure 5: **Comparison of TWASKnockoff and GIFT in simulated data with external reference panel when $h^2 = 0.02$.** We assess the performance of GIFT and TWAS-Knockoff with two feature statistics: lasso coefficients (TK-lasso) and squared z-scores (TK-squared.zscore). For each feature statistic, we consider two estimation methods (empirical estimation and bootstrap samples) of the correlation matrix of genetic elements. The $q$-value threshold is selected from 0.1 to 0.9, with the black dashed line indicating the theoretical $q$-value level. For GIFT, we applied the Benjamini-Hochberg (BH) correction to perform FDR control.

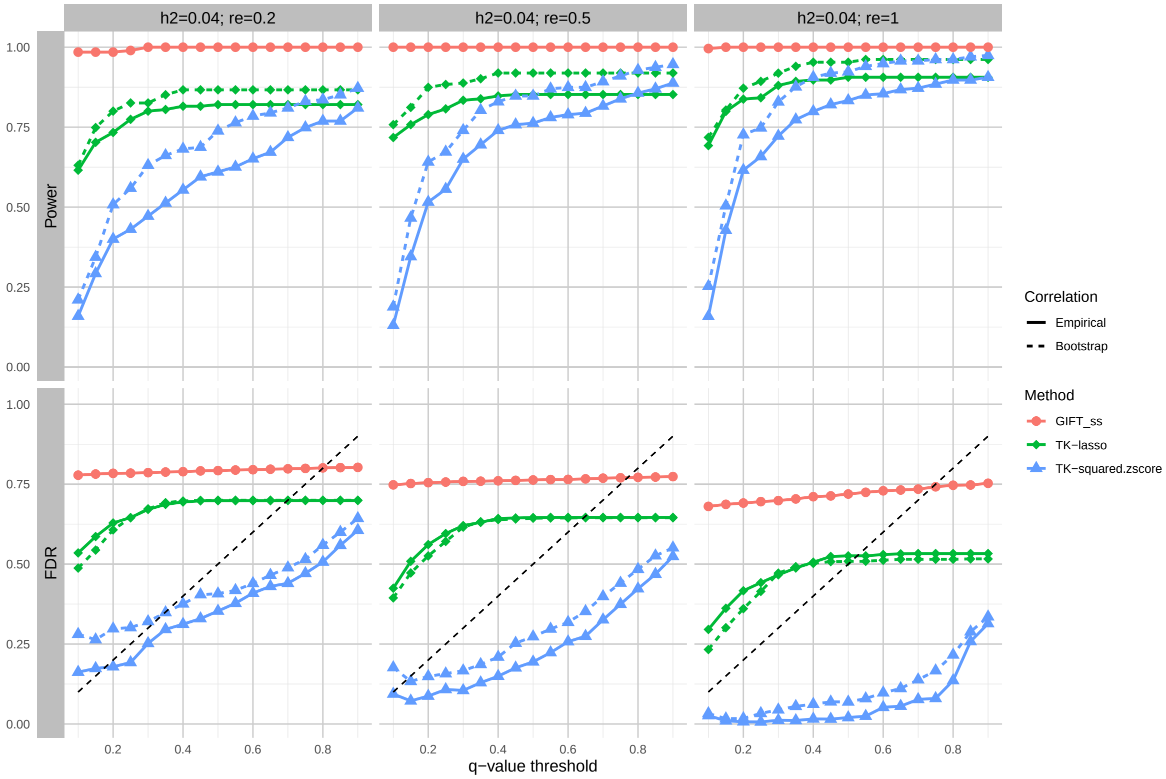


Figure 6: **Comparison of TWASKnockoff and GIFT in simulated data with external reference panel when $h^2 = 0.04$.** We assess the performance of GIFT and TWAS-Knockoff with two feature statistics: lasso coefficients (TK-lasso) and squared z-scores (TK-squared.zscore). For each feature statistic, we consider two estimation methods (empirical estimation and bootstrap samples) of the correlation matrix of genetic elements. The $q$-value threshold is selected from 0.1 to 0.9, with the black dashed line indicating the theoretical $q$-value level. For GIFT, we applied the Benjamini-Hochberg (BH) correction to perform FDR control.

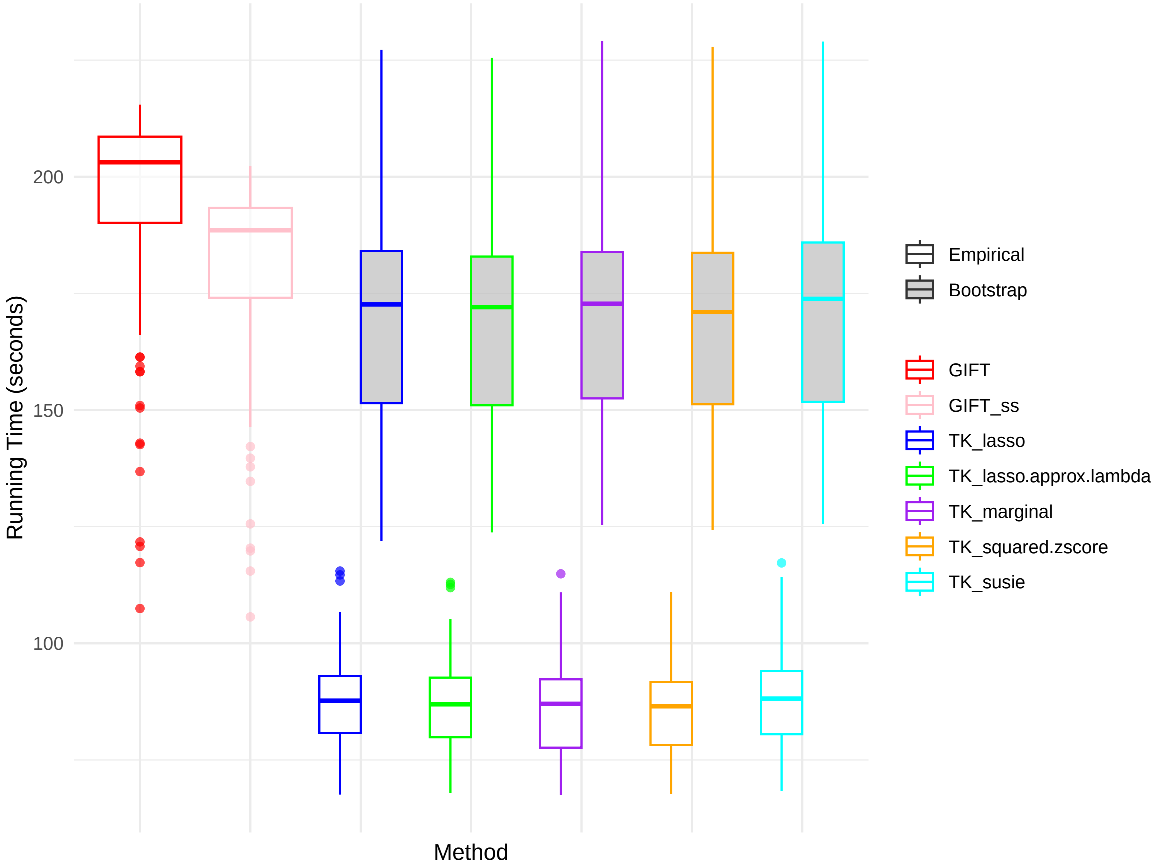


Figure 7: **Comparison of TWASKnockoff and GIFT (100 iterations) in simulated data for running time.** We assess the performance of GIFT and TWASKnockoff with five feature statistics. For each feature statistic, we consider two estimation methods (empirical estimation and bootstrap samples) of the correlation matrix of genetic elements. For GIFT, we used two versions based on individual-level data (GIFT) and summary statistics + LD matrix (GIFT_ss). In this simulation, the iteration time allowed in GIFT was set to the default number of 100, while the number of bootstrap samples for TWASKnockoff (including the estimates from the original data) was set to 10.

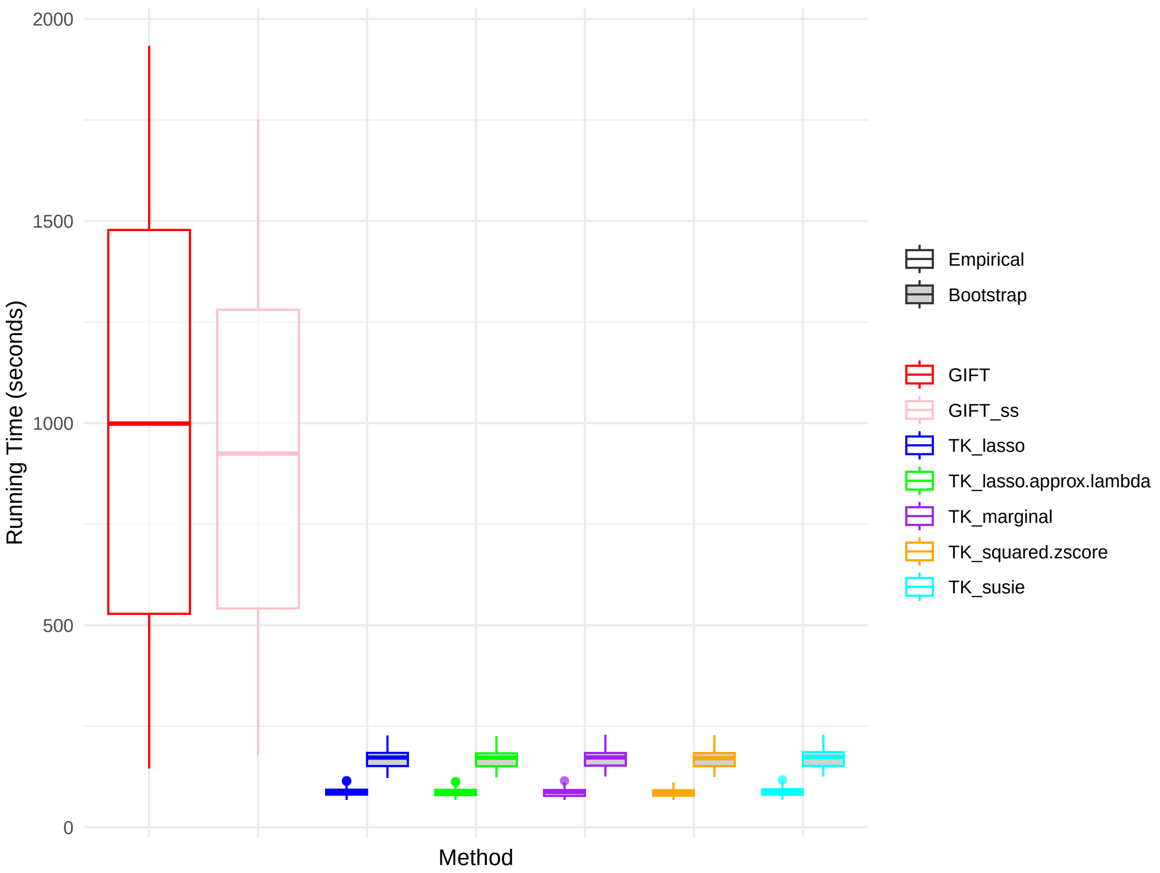


Figure 8: **Comparison of TWASKnockoff and GIFT (1,000 iterations) in simulated data for running time.** We assess the performance of GIFT and TWASKnockoff with five feature statistics. For each feature statistic, we consider two estimation methods (empirical estimation and bootstrap samples) of the correlation matrix of genetic elements. For GIFT, we used two versions based on individual-level data (GIFT) and summary statistics + LD matrix (GIFT_ss). In this simulation, the iteration time allowed in GIFT was set to 1,000, while the number of bootstrap samples for TWASKnockoff (including the estimates from the original data) was set to 10.

[2] Shaun Purcell, Benjamin Neale, Kathe Todd-Brown, Lori Thomas, Manuel AR Ferreira, David Bender, Julian Maller, Pamela Sklar, Paul IW De Bakker, Mark J Daly, et al. Plink: a tool set for whole-genome association and population-based linkage analyses. *The American journal of human genetics*, 81(3):559–575, 2007.

[3] Ken Suzuki, Konstantinos Hatzikotoulas, Lorraine Southam, Henry J Taylor, Xianyong Yin, Kim M Lorenz, Ravi Mandla, Alicia Huerta-Chagoya, Giorgio EM Melloni, Stavroula Kanoni, et al. Genetic drivers of heterogeneity in type 2 diabetes pathophysiology. *Nature*, 627(8003):347–357, 2024.
